# Unveiling molecular interactions that stabilize bacterial adhesion pili

**DOI:** 10.1101/2022.01.20.477034

**Authors:** Tobias Dahlberg, Joseph L. Baker, Esther Bullitt, Magnus Andersson

## Abstract

Adhesion pili assembled by the chaperone-usher pathway are superelastic helical filaments on the surface of bacteria, optimized for attachment to target cells. Here, we investigate the biophysical function and structural interactions that stabilize P pili from uropathogenic bacteria. Using optical tweezers we measure P pilus subunit-subunit interaction dynamics and show that pilus compliance is contour-length dependent. Atomic details of subunit-subunit interactions of pili under tension are shown using steered molecular dynamics (sMD) simulations. sMD results also indicate that the N-terminal “staple” region of P pili significantly stabilizes the helical filament structure, consistent with previous structural data, suggesting more layer-to-layer interactions could compensate for the lack of a staple in Type 1 pili. This study informs our understanding of essential structural and dynamic features of adhesion pili, supporting the hypothesis that the biophysical function of pili is niche-adapted rather than a direct consequence of genetic similarity or diversity.

## 1. Introduction

Many bacteria express micrometer long surface fibers called adhesion pili (or fimbriae), which are key virulence factors that mediate host-pathogen interactions. For Gram-negative bacteria, the most common adhesive pili class are those assembled via the classical chaperone-usher pathway (CU) [1]. P pili are an archetypal CU pilus encoded by the *pap* gene, which is significantly prevalent among strains of uropathogenic *Escherichia coli* (UPEC) that cause pyelonephritis (kidney inflammation) [2, 3]. P pili are assembled from approximately 1000 identical protein subunits (PapA) into an 8 nm thick helically wound rod with a short tip fibrillum composed of minor pilins PapF, PapE, and PapK. The fibrillum is located at the pilus distal end [4], with the adhesin protein PapG located at the very tip. This adhesin is a lectin that binds to galabiose-containing glycosphingolipids [5].

To assemble a P pilus all subunits (pilins) are transported from the inner membrane through the periplasm via the general secretory pathway [1]. During their transport to the outer membrane by the PapC usher [6], the pilins are folded and stabilized by the periplasmic chaperone PapD via donor-strand complementation (DSC) [7]. At the outer membrane each pilin subunit is transferred from the chaperone to the usher, where it binds to the linear polymer of previously assembled subunits via donor strand exchange (DSE). As the polymer is assembled, subunits are translocated to the cell surface through the usher’s central pore [1]. After exiting the pore, the PapA polymer forms a quaternary helical surface filament of 3.28 subunits per turn with a pitch of 25.2 Å and a diameter of 81 Å [8, 9]. Each subunit *n* in the helical filament is bound through hydrophobic and weak hydrophilic interactions with ten other subunits, that is, with five preceding (−5 to -1) and five succeeding (+1 to +5) subunits, forming a large network of interactions (Figure 1). The extension of this network to such distant subunits in P pili (*n* to *n*-4 and *n*-5) is primarily due to the staple that is comprised of the first seven residues of the PapA N-terminus. It is therefore hypothesized that the staple has an extensive stabilizing role despite the fact that the staple region might not be essential in rod formation [9]. Interestingly, so far, the staple region has only been found on P pili and is missing in closely related UPEC pili such as Type 1. Also, this staple region is missing in the archetypal CFA/I pili expressed by enterotoxigenic *Escherichia coli* (ETEC) [10].

**Figure 1:**
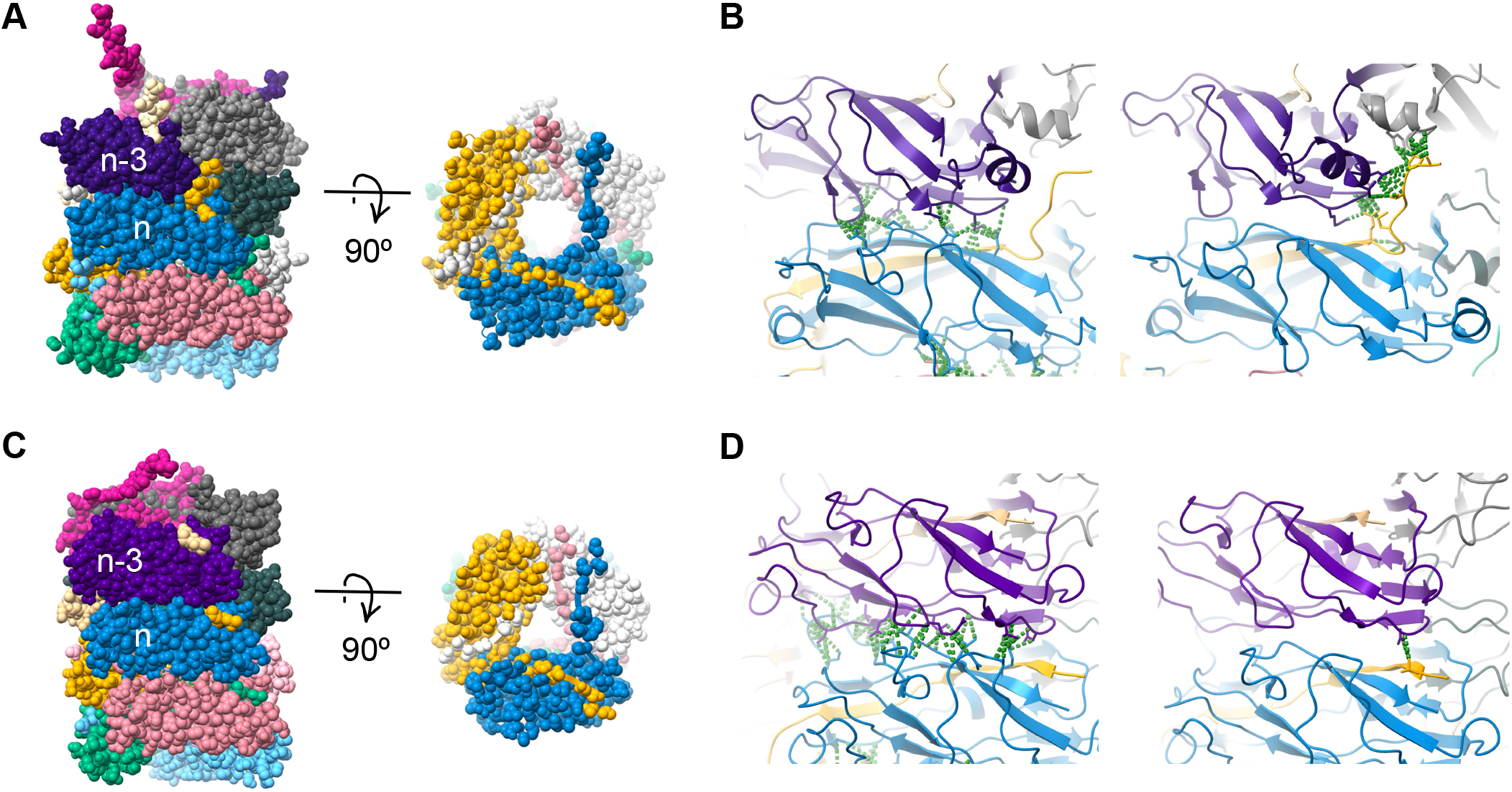
The staple region of P pili provides specialized subunit-subunit interactions. **A**) A surface view of P pili with subunits individually colored; the N-terminal extension of the *n*+1 yellow subunit is inserted into a groove in the *n*^th^ subunit and has a staple that forms contacts with subunits *n*-3 and *n*-4. **B** Contacts between P pili subunits are shown in green between subunits *n* and *n*-3 (left, layer-to-layer interactions) and between subunit *n*+1 to *n*-3 and *n*-4 (right, staple interactions). **C** Type 1 pilus structure, for comparison with panel A. **D** Type 1 pili have increased *n* to *n*-3 interactions (left) and only 1 contact of *n*+1 to *n*-3, as there is no staple region; *cf* panel B.

Despite the large interaction network of subunits, the bulk of the surface contact area is between the *n*^th^ subunit and -3 and +3 (Figure 1) [9]. Not surprisingly, the magnitude of the surface contact area is closely related to the force needed to uncoil the quaternary structure of helical pili. For example, from cryo-EM structures and force-extension experiments it has been shown that PapA subunits have a buried surface area between *n* and *n* + 3 of 1453 Å^2^ and P pili require 28 pN of force to uncoil, whereas FimA in Type 1 pili and CfaB in CFA/I pili have 1615 Å^2^ and 1087 Å^2^ and require 30 pN and 7 pN of force to uncoil, respectively [9–12]. The experimentally measured forces needed to uncoil pili are of the same magnitude as the fluid flow forces expected to be present in the *in vivo* environmental niches of bacterial cells [13]. The ability of pili to uncoil by breaking sequential layer-to-layer interactions reduces the shear forces on the pili, and is thereby expected to aid sustained bacteria adhesion under fluid flow [14]. This hypothesis is supported by *in vivo* data in which bacteria expressing mutant Type 1 pili, with reduced surface contact area between layer-to-layer subunits, had a significantly reduced ability to cause intestinal colonization and bladder infection in mice [15].

Thus, the specific biophysical properties of pili are crucial for bacterial adhesion. Over the years, numerous helical filament pilus types have been characterized. These measurements revealed that pili are elastic and superelastic filaments and that they can be completely uncoiled into a linear fiber under force. When the force is removed, the structure can regain its helical quaternary shape [12, 16–24]. Although several studies have been performed and models developed to explain the superelasticity behaviour, there are still several open research questions that need attention. For example, does the staple found in P pili subunits provide a stabilizing role when tensile stress is applied, and are there other possible functions? How do subunits re-orientate when unbound? How much longer does a pilus get upon extension? This number varies in the literature. While the extension response of a pilus is well described by biophysical models, these models treat subunits as individual blocks connected by springs and do not consider individual molecular interactions. Therefore, questions about the contact network between pilins are still unanswered. For example: What contacts are most important for pilus stability? How fast are the transitions between the subunits’ bound and unbound states? Are there intermediate or alternative states that the subunits can visit during binding and unbinding? To answer these research questions, we combined optical tweezers (OT) force experiments and molecular dynamics (MD) simulations to unveil the molecular interactions that stabilize chaperone-usher pili and their role for pilus mechanical and kinetic properties.

## 2. Methods

### 2.1. Bacterial strains and growth conditions

We used the *E. coli* strain HB101 as host strain for the plasmid pHMG93 to express UPEC-related P pili [25–27]. We cultured the bacteria on trypticase soy agar at 37 ^°^C for 24 h.

### 2.2. Optical tweezers force measurements

To apply strain and thereby extend a pilus and track its length fluctuations we used an in-house built OT setup. The OT setup uses an inverted microscope (Olympus IX71, Olympus, Japan) as a base. To image the sample and form the trap we use a water immersion objective (model: UPlanSApo60XWIR 60X N.A. = 1.2; Olympus, Japan) and a 1920 × 1440 pixel CMOS camera (model: C11440-10C, Hamamatsu) [28]. We minimized the amount of noise in the OT setup by using Allan variance analysis [29], and we used an active Power Spectrum method to calibrate the trap [30]. During calibration we oscillated the bead at 32 Hz with an amplitude of 100 nm. We sampled the microbead position at 131,072 Hz and averaging 32 consecutive data sets acquired for 0.25 s each. In general, the trap stiffness was 370 pN/*µ*m. An example of a power spectrum with a corresponding fit from our instrument is shown in Figure S1.

To extend a pilus we first attached a bacterium to a poly-l-lysine coated immobilized microsphere, and then trapped a bead with our optical trap. The trapped bead was moved in proximity to the bacterium to attach a pilus. Thereafter, we moved the piezo stage at a speed of 50 nm/s, below steady-state, to apply tensile fore. To measure the kinetic of subunit opening we uncoiled 150 nm of the rod and then kept the piezo stage and trap stationary. We sampled the force and position of the bead at 100 kHz with an anti-aliasing filter set to 50 kHz. We recorded 5 data series of 30 s each for a total of 2.5 min of data per pili. We show a schematic with details of the setup and more information of the measurement procedure in section S0 and Figure S2 in Supporting Material.

### 2.3. Force Data Analysis

We averaged the time series data to an effective sampling rate of 500 Hz to remove thermal noise and increase spatial resolution. As we measured using a constant trap position, instead of constant force, the measured displacements of the bead did not reflect the true length changes of the pili. To correct for this effect we did compliance corrections for each data set using the stiffness of the pilus. For details on how We retrieved the stiffness from variance of the bead fluctuations and did the compliance correction, see section S1 in Supporting Material. As the bead is connected to the trap and pilus in parallel their stiffness is additive, so that difference of the stiffness when attached to the pilus and that for the bead in the trap alone gives an estimate of the stiffness of the pilus. With this estimated stiffness we generally determined a correction factor of approximately 1.5, indicating a stiffness of approximately 700 pN/um. We then analyzed the corrected data by taking a sliding window histogram with a window length of approximately 1s of the data to retrieve the distance between states.

### 2.4. Molecular dynamics simulations

The initial structure of the *E. coli* P pilus was obtained from PDB entry 5FLU [9]. Simulations were carried out for a 7mer segment of the filament following a protocol very similar to [11], but with some changes made related to the larger system size simulated here. The program Amber20 was used to perform all simulations for this work [31]. Steered molecular dynamics simulations were carried out at constant velocities of 1 Å/ns and 5 Å/ns with the staple region of the PapA N-terminal extension present, and also with amino acids in the staple region removed. In each case (each pulling speed and with/without the staple region) five simulations were performed. Filament extension in the steered molecular dynamics simulations was directed along the z-axis which is aligned with the filament axis. For additional details, please see the description in the Supporting Material.

## 3. Results and Discussion

### 3.1. Force-extension experiments unveil the kinetics of subunit opening and a contour length-dependent compliance

To study the layer-to-layer interactions that stabilize the P pilus rod and the consequences of bond breakage for pili mechanics, we used OT. OT measurements, and to some extent atomic force measurements, have shown that helix-like pili often exhibit two or three different modes of elongation, which show up as distinct regions in their force-extension responses [18, 32]. These regions are well described by elastic and entropic elasticity (region I), superelasticity (region II), and a combination of elastic and entropic elasticity with a phase transition (III) (Figure 2A) [11, 33]. In Figure 2A, we can see that the initial force response (region I) of a P pilus continues linearly until it reaches a threshold force of 28 pN. At 28 pN, we enter region II, where layer-to-layer interactions break, resulting in a sequential uncoiling of the P pilus rod. Due to the tight native packing of the pili, 3.28 subunits per turn [8], the sequential uncoiling causes the pilus to elongate [18], as illustrated in Figure 2B (panel I), keeping the force experienced by the pilus close to constant.

**Figure 2:**
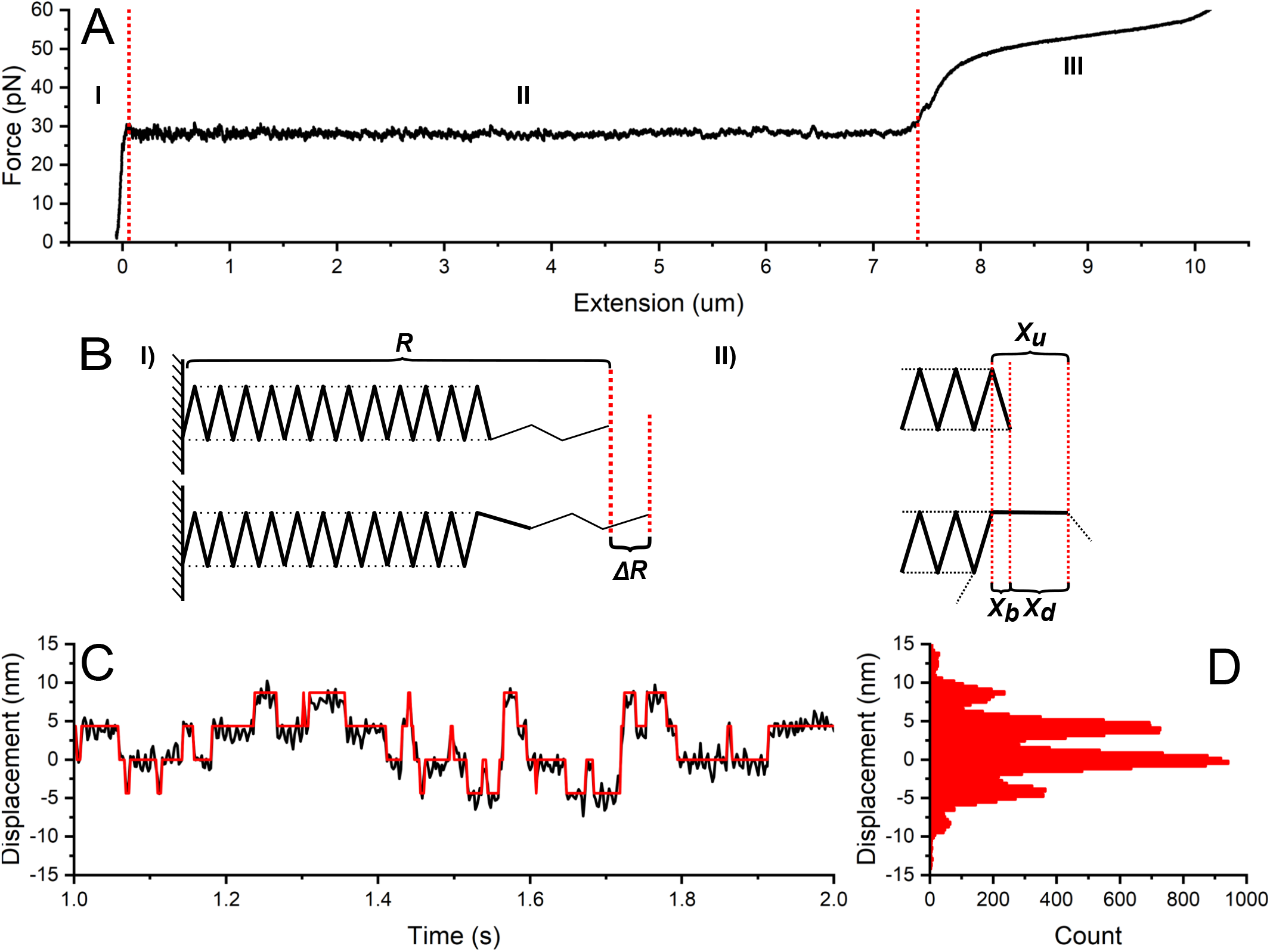
P pili uncoil under force in a step-wise manner. **A**) An example of a force-extension curve of a P pilus. The different regions are marked with I, II, and II. Boundaries between the regions are demarcated by dashed red lines. **B** I) Illustration showing the helical structure of a P pilus with a total end-to-end distance *R* made out of many bound subunits (solid lines) in which layer-to-layer interactions (dashed lines) can break and reform, causing an end-to-end distance change Δ*R*. II) Zoomed-in view of the layer-layer-interaction breaking showing the contour length contribution of a bound (folded) subunit *X*_*b*_, the contribution of an unbound subunit *X*_*u*_, where *X*_*d*_ is the difference in pilus contour length after a subunit uncoils **C** An excerpt from a time-series showing the displacement of the trapped microbead. The displacement states are shown in red. **D** Displacement histogram of the time-series data showing discrete peaks due to the changing number of bound and unbound subunits.

However, although the force in region II appears constant, it is rapidly fluctuating due to thermal energy that is breaking and reforming the layer-to-layer interactions that bind the subunits in a coiled formation. That is, subunits randomly shift between their bound and unbound state. As such, we would expect a pilus held at a constant force in region II to undergo random length changes over time. To study these length changes, we extended the pilus and uncoiled it roughly 150 nm into region II, and then held the bead stationary. We stopped at 150 nm extension since pili get softer as they uncoil, reducing temporal and spatial resolution in our measurements; a data set showing this softening and the change in the distribution of fluctuations of a pilus measured at two positions along region II, is shown in Figure S3. Therefore we measured the random length changes of pili that were mostly in the helical, coiled arrangement with only a small region of the pilus uncoiled. Using this procedure, we obtained a time series of small length changes in the trapped pilus. For these displacements to reflect the pilus’s end-to-end length changes (*R* in Figure 2B), we included a compliance correction. This compliance correction accounts for the effects of the finite trap stiffness; see section S1 in Supporting Material for details on the compliance correction. We show an example of a two-second section from a time series in Figure 2C. These data show clear random transitions between discrete displacement states corresponding to the varying number of subunits in the bound or unbound state.

To analyze these state transitions in detail, we plotted a histogram of the measured length fluctuations, as shown in Figure 2D. This histogram shows that the pilus mostly visits six discrete states, corresponding to six subunits randomly being in the bound or unbound state. Further, we find that the spacing between these states is 4.2±0.2 nm (n=10 000 transitions, 10 bacteria, 2 biological replicates). This value corresponds to the change in end-to-end length of the pilus when a single subunit switches from the bound state to the unbound state, shown as Δ*R* in Figure 2B (panel I). However, as the uncoiled region of the pilus is highly flexible (2 nm thick), thermal fluctuations will contract it due to conformational entropy. Thus, the mean contribution of a subunit to the total end-to-end distance will be less than its contribution to the length of the pilus backbone, the contour length. For these distance definitions, see Figure S4. To relate the change in contour length to the measured end-to-end distance change, we created an entropic elasticity model for a partially uncoiled pilus, as described in section S1 of Supporting Material. Our entropic elasticity model combines a worm-like chain model and a free-jointed-chain model that describes the coiled and uncoiled part, respectively. Comparing this model to a previous model [33] shows a significantly better agreement with experimentally measured pilus stiffness, Figure S5. Using this new stiffness model, we find that the increased contour length contributed by a single subunit going from the bound to the unbound state is 4.35±0.2 nm, corresponding to the contour length increase of the pilus after a subunit unbinds, *X*_*d*_ in Figure 2B (panel II).

We can relate this value, *X*_*d*_, to the length contributed by a single subunit bound within the coiled region, *X*_*u*_, through the relationship *X*_*u*_ = *X*_*d*_ + *X*_*b*_. The contour length contribution *X*_*b*_ of a bound subunit is known from the molecular structure and is 0.754 nm [34]. Thus, we can estimate *X*_*u*_ to be ∼5.1 nm (4.35+0.75 nm). This value is close to the ∼5 nm length of a PapA subunit [35]. Thus, we can infer that the donor strand, which acts as a hinge between the subunits, does not restrict subunit movement during uncoiling, allowing the pilus to become fully linearized.

Knowing this, we can also estimate the pilus elongation ratio, the ratio between its coiled and uncoiled length. We determine this by taking the ratio of a subunit’s bound and unbound length. Thus, the elongation ratio is given by (5.1-0.75)/0.75 = 5.8, which means that a P pilus gets roughly 6 times longer after uncoiling. This elongation ratio lands in the middle of the previously reported values for P pili, which reported 4-7 times elongation [8, 16, 18]. Further, it is worth noting that our value of *X*_*d*_ is larger than previously reported for P pili, where a change in end-to-end distance was assessed to 3.5 nm [18]. Most likely, this difference originates from the simple geometrical and kinetic models used in that work to describe the uncoiling of pili.

In addition, we see that the histogram in Figure 2D indicates no significant intermediate states between the bound and unbound states in our force experiments. To ensure that we did not miss any fast transitions due to a low sampling rate, we increased our sampling rate from 500 Hz to 100 kHz, the limit of our system. However, even sampling at this speed did not indicate additional intermediate steps. Therefore, if there are intermediate states between the bound and unbound states of a subunit, they must exist on a shorter time scale than the millisecond response time of our setup. Further, it is worth noting that we do not see any indication of P pili recoiling at two force levels, as is seen in some force curves of Type 1 and S_*II*_ pili [19, 20, 36]. In particular, for Type 1 pili, this observation might reflect that the rod can adopt two different folding configurations of the quaternary structures, as observed when comparing Type 1 pili assembled either *in vitro* or *in vivo* [37, 38].

To conclude, our force measurement results indicate that the process of going from the bound to unbound state is faster than 10 msec, so we cannot resolve any intermediate steps. Therefore we cannot assess experimentally the detailed changes in subunit-subunit interactions that occur during uncoiling. However, a model from cryo-EM data shows multiple stabilizing interactions between subunits in the coiled configuration [9]. Thus, we turned to MD simulations to investigate these interactions more closely and validate our results for the length change assessment upon uncoiling.

### 3.2. MD reveals mechanistic details of pilus uncoiling

We carried out steered molecular dynamics (sMD) simulations of a 7mer segment of P pili. By simulating a 7mer filament, we can investigate the process of bond-opening as the pilus is elongated in the direction of the filament axis. The sMD simulations were carried out at two different, constant pulling speeds, 5 Å/ns and 1 Å/ns, with five runs for each pulling speed. We also simulated a version of the 7mer system with the staple region removed (specifically, without amino acids 1-5 present) to investigate in our model the contribution of staple residues to the force required to extend the 7mer. Here we describe in detail the results of one run of the 1 Å/ns simulations with the staple, which displayed very similar features compared to all other runs at both pulling speeds. Comparisons to the simulations carried out without the staple are described below. Details of the sMD protocol are described in Methods, and data for all other runs of both systems are included in the Supporting Material.

To demonstrate the major reproducible features observed during filament extension in Figure 3A we show all five runs of the 1 Å/ns sMD simulation force-extension curves plotted together. Each of the first four peaks corresponds to a particular subunit that is being pulled away from the 7mer filament. For example, peak I represents the subunit at the pilus tip (subunit 1) being pulled away from the filament, while peaks II-IV represent the subsequent breaking of layer-to-layer interactions for the second, third, and fourth subunits in the filament, respectively. Since we placed positional restraints on the bottom three subunits of the filament to provide a stable filament rod against which pulling occurs, we did not observe force peaks corresponding to extension of the bottom three subunits. Instead, because the base was restrained, as we continued to pull on the filament at constant speed, a breakage event occurs somewhere in the filament once the pilus was fully extended (e.g., a donated beta strand being pulled out from a subunit). Peak V represents this breakage event, and the same behavior was observed in simulations of the P pilus 3mer [11]. Force-extension curves for the additional simulations of the 7mer system at a pulling speed of 5 Å/ns, as well as the 7mer system without the staple at both pulling speeds, are included in Supporting Material Figure S6. These data demonstrate that we observed similar features across both pulling speeds and across all simulation runs for the wild-type system. Finally, we note that filament breakage events can occur at various points along the filament in our simulations (e.g., the final panel in Figure 3E). Movies of each of the sMD trajectories are found in the Supporting Material (see Movies S1-S4).

**Figure 3:**
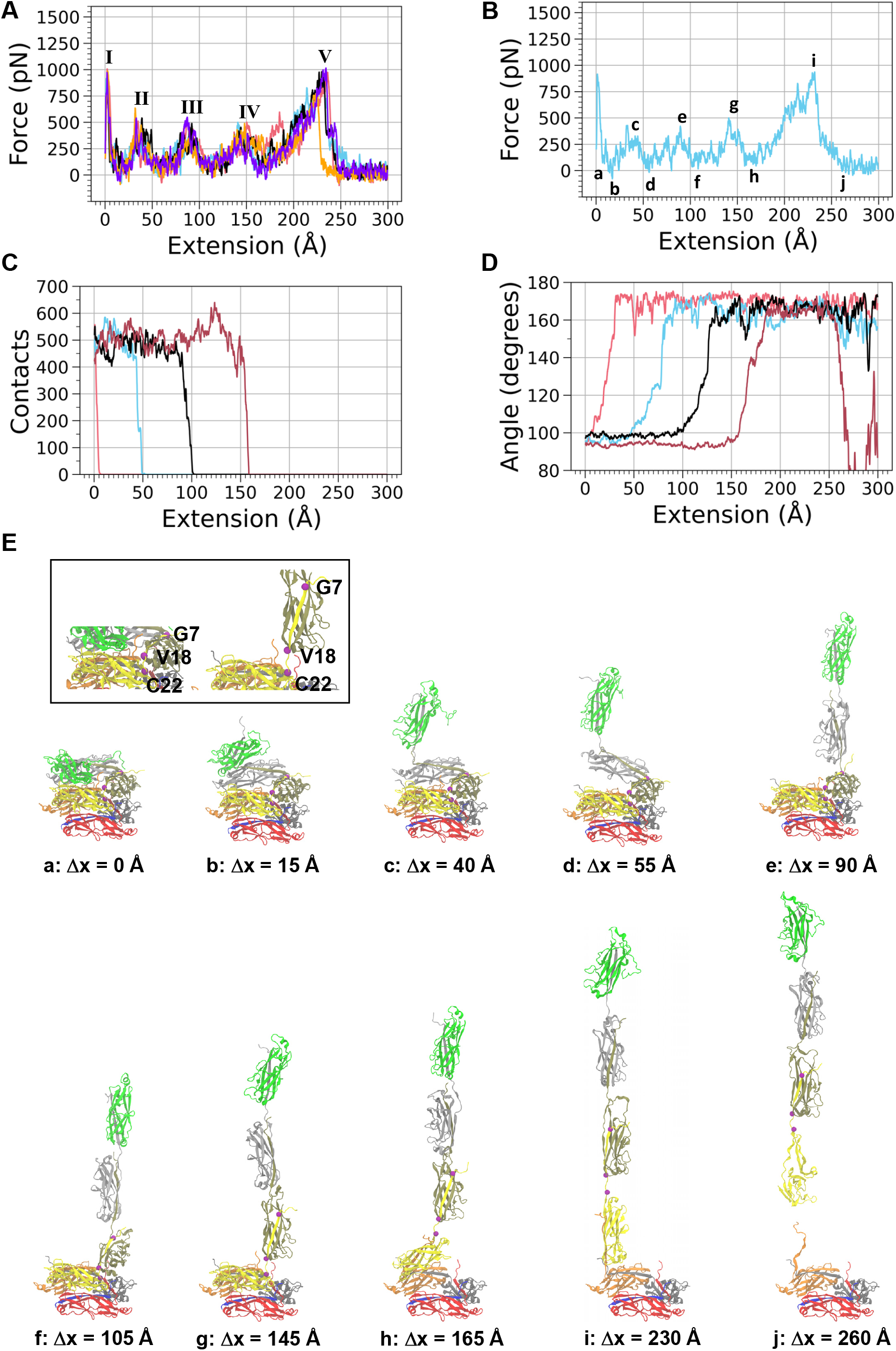
Force, contacts, and subunit interactions as P pili uncoil under force via steered molecular dynamics. **A**) Force versus extension curves for all five of the v = 1 Å/ns steered molecular dynamics simulations with the staple present (pink = run 1, blue = run 2, black = run 3, orange = run 4, purple = run 5). Roman numerals I-IV identify the peak force at the point that a subunit was unwound from the filament, and Roman numeral V represents the force right as the 7mer was severed between two subunits (e.g., through the removal of a donated beta strand). The force curve data are a 1 ns running average. **B**) The force versus extension data for run 2. Lowercase letters along the curve indicate which image in panel E of this figure corresponds to that degree of extension. **C**) Total number of native plus non-native contacts between subunits 1 (tip subunit) and 4 (pink curve), 2 and 5 (blue), 3 and 6 (black), and 4 and 7 (dark red). For this analysis, a contact is defined as two atoms in a pair of subunits coming within a 4 Å cutoff distance of one another. The contact data are a 1 ns running average. **D**) Angle between residues Gly 7, Val 18, and Cys 22 (purple spheres in panel E). The curves correspond to the alignment of subunit 1 (pink curve), subunit 2 (blue curve), subunit 3 (black curve) and subunit 4 (dark red curve) with the filament axis. When the angle is 90 °, the subunit is perpendicular to the filament axis and when the angle is 180 °, the subunit is aligned with the filament axis. The angle data are a 1 ns running average. **E**) Snapshots of the sMD simulation with letters that correspond to the labels in panel B; purple spheres represent the positions of Gly 7, Val 18, and Cys 22.

Specific features observed in the simulations are illustrated by a single run from the 1 Å/ns simulations in 3B, C, D, and E. The lowercase letters at various points along the force-extension curve in Figure 3B correspond to the images in Figure 3E that depict representative snapshots from an sMD trajectory. The first peak (immediately after **a**) and the peaks at **c, e**, and **g** correspond to bond-breaking events as subunits were pulled away from the filament rod. We note that these peaks coincide with the approximate extension at which the total number of contacts between subunits *n* and *n*+3 (i.e., the subunit pairs which make lateral layer-to-layer interactions) began to fall rapidly towards zero (Figure 3C). Similarly, the valleys at **b, d, f**, and **h** correspond to approximate extension lengths at which the contacts between a pair of *n* and *n*+3 subunits had fallen identically to zero (Figure 3C); see Supporting Material Figures S7-S10 for all plots of contacts versus filament extension, including systems both with and without the staple.

We observed that at the point where the contacts reached zero between *n* and *n*+3 subunits, the angle of rotation of the newly freed subunit began to rapidly change with respect to the filament axis (Figure 3D, and Figures S11-S14). We used the angle made by three amino acids (Gly 7, Val 18, and Cys 22), which are shown in the inset image of Figure 3E, as a proxy for the change in the angle of the subunit with respect to the filament axis. The angle between those amino acids started at approximately 95 degrees and rotated to become nearly linear, corresponding to the linearization of the subunits as they were extended away from the filament rod due to the applied force (Figure 3E).

Taking Figure 3B, C, D together as a representative example of filament extension, we can infer how the bond-breaking process generally occurs as the subunits are pulled away from the filament in the P pilus. For example, we can consider the points **b, c**, and **d** in Figure 3B which demarcate a full force peak. At point **b** the force was beginning to increase to a maximum as the applied force was not yet strong enough to break all of the contacts between the second subunit (light grey) and the fifth subunit (orange) in the filament. Once the peak force was achieved at point **c** contacts began to rupture (Figure 3C, blue curve) and then as the number of contacts dropped rapidly to zero the force decreased between points **c** and **d**. Once the force had reached a minimum at point **d** the second subunit was free to rotate, becoming parallel to the filament axis (Figure 3D, blue curve). The force again began to increase between points **d** and **e** and the contacts between the next pair of *n*/*n*+3 subunits began to rupture. This process was then repeated for each of the subunits in the filament. We therefore infer that, in a simulation of a much longer length of filament, we would continue to observe this sequential bond-breaking pattern, connecting the applied force, contact breakage, and subunit rotation.

Similar to the 4.3 nm bond opening length observed in OT experiments, we note that the extension over which a subunit rotation occurs, as observed in Figure 3D, is also in this range. For example, for the pink curve (corresponding to the rotation of the terminal subunit) the rotation started at ∼0 Å and stabilized at a nearly straight angle by ∼35 Å extension. For the next three curves (blue, black, red), which correspond to rotation of the second (grey), third (goldenrod), and fourth (yellow) subunits respectively, the rotation from low angle to high angle occurred over approximately 45 Å, 40 Å, and 45 Å, respectively. Therefore, each bond-opening rotation of a subunit contributes ∼35 - 45 Å of length toward extending the filament, in close agreement with the experimental data. Additional elongation outside of the bond-opening events is therefore related to additional flexibility of other components of the subunits as, for example, observed upon unfolding of an alpha helix within a subunit [11].

### 3.3. The staple region impacts quaternary stability and subunit-subunit interactions

As discussed above, we were also interested in investigating the effects of the staple amino acids on the response of the P pilus to force. It has been previously reported that these amino acids contribute a significant amount of buried surface area to the interface between subunits in the pilus filament [9], and therefore these amino acids might provide additional stability against force. To ascertain the importance of the staple for pilus subunit-subunit interactions when pili are subjected to force, we deleted the first five amino acids at the N-termimus of each subunit and repeated the sMD simulations.

For the simulations in which the staple was removed, we note that an overall lower average level of force was required to unwind each subunit from the helical filament, compared to simulations in which the staple was present (Figure 4A, 1 Å/ns). This lower force for unwinding suggests that the staple amino acids provide stabilizing interactions to the pilus filament. The averaged force data for the systems both with and without the staple at a pulling speed of 5 Å/ns are shown in Figure S6D, which shows the same trend in the overall average level of force for the two systems as is observed in Figure 4A.

**Figure 4:**
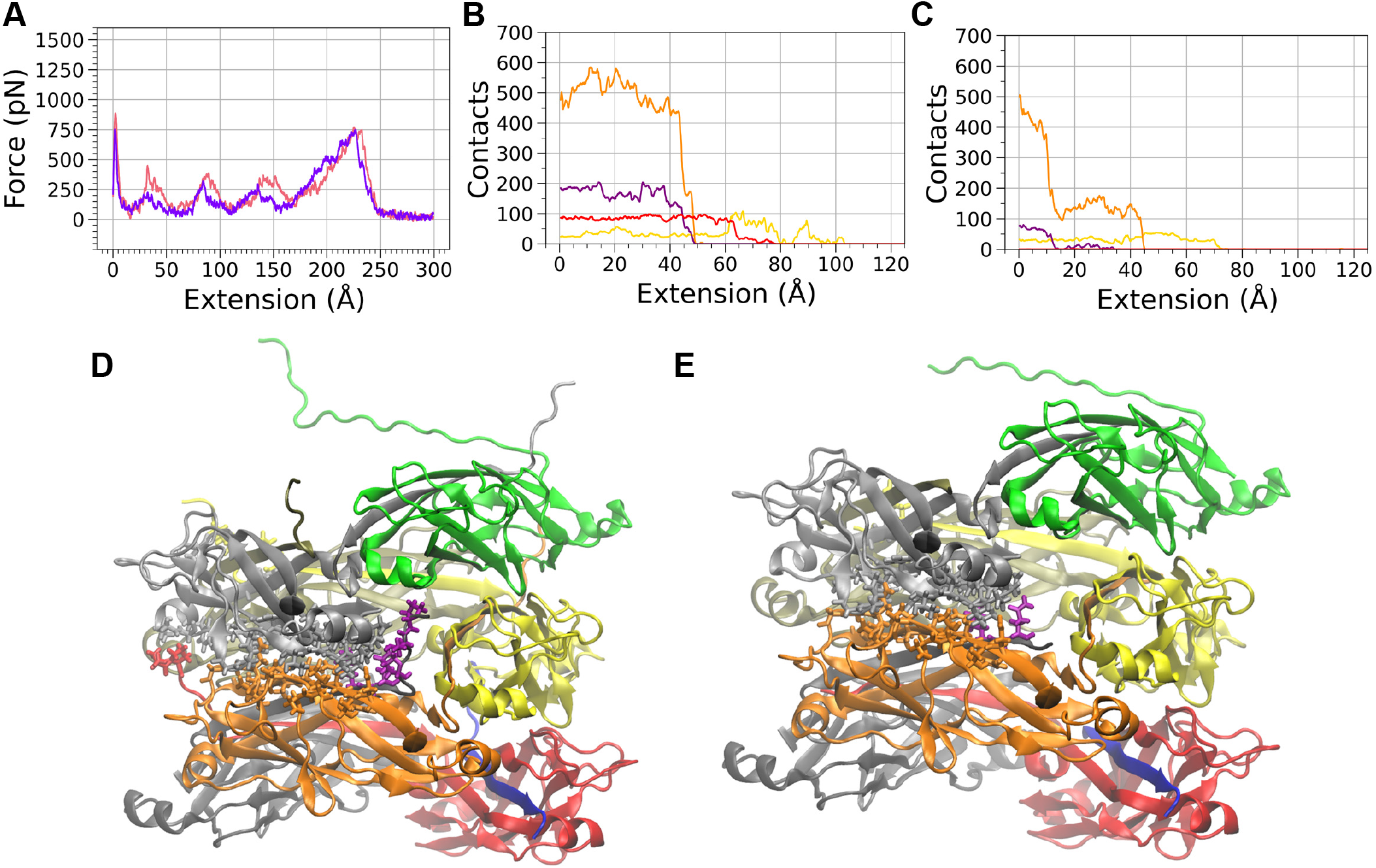
The staple region provides stability to P pili under force. **A**) Force versus extension curves for the v = 1 Å/ns steered molecular dynamics simulations averaged over five separate runs for the system including the staple (pink) and with the staple residues removed (purple); the data are a 1 ns running average. **B**) Total number of contacts between subunits 2 and 4 (yellow), 2 and 5 (orange), 2 and 6 (purple), and 2 and 7 (red), for run 2. Contacts data are a 1 ns running average. They were calculated as described in Figure 3C. Data are for the simulations with the staple. **C**) The same contact analysis as shown in Panel B, except for run 2 of the v = 1 Å/ns simulation *without* the staple. Panels **D**) and **E**) show images of the 7mer with subunit 2 atoms (light grey subunit) initially within 4 Å of subunits 4 (yellow), 5 (orange), 6 (dark grey), and 7 (red). The corresponding residues of subunits 4, 5, 6 and 7 that are within 4 Å of subunit 2 are also shown. For the residues on subunit 6, atoms are colored in purple for clarity. Colors of the lines in panels B and C are the same as the subunit coloring in panels D and E, except the contacts between subunit 2 (light grey) and subunit 6 (dark grey) are also drawn in purple for clarity.

We also observed that in simulations with the staple amino acids removed, contacts between subunits were eliminated over a smaller range of filament extension for the majority of runs, compared to simulations in which the staple was present (compare Figures S7 and S8, as well as Figures S9 and S10). To understand these trends in the contact breakage more fully, we also analyzed the contacts formed between the second subunit (light grey subunit in Figure 3) with each of the four surrounding subunits (the 4th, 5th, 6th, and 7th subunits in the 7mer filament). The data for this analysis of the sMD simulations at 1 Å/ns is shown in Figure 4B, C (run 2), and all of the data for 1 Å/ns simulations are shown in Figure S15. By analyzing the contacts separately instead of in aggregate, we can observe more directly how the interactions between a subunit being pulled off of the filament rod and other subunits around it were disrupted. As expected, the largest number of contacts between subunits occurred for the layer-to-layer *n*/*n*+3 interactions (orange curves). This is true for both the simulations with and without the staple. However, in simulations without the staple, the number of layer-to-layer contacts generally dropped more quickly than when the staple was present; with the staple present, the layer-to-layer contacts between subunits *n*/*n*+3 began to drop roughly simultaneously with the contacts between the 2nd and 6th subunit. Contacts formed between these two subunits primarily came from the staple region of the 6th subunit, as seen in the figures. We note that some interactions of the 2nd subunit also occurred with the staple of the 7th subunit (red curves), and were therefore absent without the staple. Contacts between the 2nd subunit and the 4th subunit were the last to be completely broken in each of the simulations, which occurred as the filament was straightened and pulled away from the filament rod (e.g., Figure 3E).

Finally, to define more specifically some of the important contacts that occur between amino acids in the P pilus, both in the presence and the absence of the staple amino acids, we analyzed data from our 100 ns equilibrium simulations to determine the most stable interactions. As seen in Tables S1-S3, we observed that some interactions found to be highly stable in the 100 ns equilibrium simulations were also noted in the P pilus structure by Hospenthal et al [9]. For example, a particularly large number of stable interactions were observed between Lys 125, Asp 126, and His 132 with other amino acids in our simulations. This finding is consistent with the importance of those amino acids for P pilus stability, as indicated by the mutation experiments in Hospenthal et al. demonstrating that their mutation led to diminished rod stability. Interestingly, in Hospenthal et al. the mutation of Thr 3 (slightly polar) to an arginine (positively charged) produced helical pili comparable to wild-type, and in our simulations without the staple, while we observed a decreased unwinding force for the P pilus, sMD elongation of the system is mechanistically similar to wild type. Together, the data suggest that Thr 3 is not a critical residue for pilus dynamics. Other changes that occur when the staple is removed include a small number of new interactions that are only present in the equilibrium simulation without the staple (Table S4). It is possible that these new interactions could occur after removal of steric hindrance by the staple, producing a slightly stronger pilin-pilin interaction.

Mutation experiments in combination with OT force measurements, similar to the approach by Spaulding et al. [15], could test the presence and role of a modified staple region in P pili. This would allow for an interesting comparison of P and Type 1 pili since Type 1 lacks the staple. It would then be possible to determine the contribution of the staple to the observed ∼50 times higher bond opening rate for P compared to Type 1 pili [19], and explore a possible role for the staple in guiding subunits into their bound state during recoiling. This is plausible since the staple increases the “reach” of the layer-to-layer interactions, which could increase the probability of an unbound subunit finding its way back to the coiled subunits.

In summary, we investigated the biophysical properties of P pili and the structural interactions that stabilize this representative of the helical class of the chaperone-usher adhesion pili family. Using high-resolution force measuring optical tweezers, we unveil contour-length dependent compliance of helical pili: a pilus starts very stiff (700 pN/µm) but softens significantly during uncoiling (100 pN/µm). We find that this biophysical property is well described by a coupled WLC and FJC model. Further, we assessed the kinetics and orientation change of subunits unbinding from the rod. We found that at steadystate (the plateau force), at most six subunits are in the unbound state. When a single subunit switches from the bound state to the unbound state, the end-to-end length of the pilus increases 4.35 nm. A pilus can thus extend about six times its coiled length. Taken together, these experimental results imply that the donor strand, which acts as a hinge between the subunits, does not restrict subunit movement during uncoiling, allowing the pilus to become fully linearized.

We verified these findings, investigated the bond-breaking process, and investigated the molecular interactions that stabilize the helical structure using 7mer filament sMD simulations. With a 7mer system our simulations provide the first molecular scale view of P pilus uncoiling at an atomistic level of detail, in a system large enough to include layer-to-layer interactions, aiding our interpretation of experimental force measurements. This detailed view of how interactions break and the virtual removal of the staple region in sMD simulations provided valuable information about the role of the staple for P pilus stability. From our results we infer that the staple region significantly helps stabilize the helical rod structure. This study investigated essential features found in the helical class of uropathogenic adhesion pili and adds momentum to the observation that the biophysical properties and the functioning of pili are niche adapted.

## 4. Acknowledgements

This work was supported by the Swedish Research Council (2019-04016) to M.A. The authors acknowledge the facilities and technical assistance of the Umeå Core Facility for Electron Microscopy (UCEM) at the Chemical Biological Centre (KBC), Umeå University, a part of the National Microscopy Infrastructure NMI (VR-RFI 2016-00968). J.L.B. acknowledges support under NSF Grant MCB-1817670. J.L.B. also acknowledges use of the Electronic Laboratory for Science and Analysis (ELSA) high-performance computing cluster at The College of New Jersey for conducting the simulations reported in this paper. This cluster is funded, in part, by the NSF under Grants OAC-1826915 and OAC-1828163. This work was supported by the NIH, R21-AI-156236 to EB.

