## Supporting information for "Unveiling molecular interactions that stabilize bacterial adhesion pili"

[a]Department of Physics, Umeå University, Umeå, 90187, Sweden;
[b]Department of Chemistry, The College of New Jersey, Ewing, NJ, 08628, USA;
[c]Department of Physiology & Biophysics, [c]Boston University School of Medicine, Boston, MA, 02118, USA
[d]Umeå Centre for Microbial Research (UCMR), Umeå, Sweden
[e]Contributed equal to the work

### Supporting Material content

**S0 Optical tweezers instrumentation and sample preparation** **2**

**S1 Supporting force spectroscopy information and pilus modeling** **5**
Measuring stiffness of an uncoiled pilus 5
Compliance correction 6
Stiffness model for an uncoiling pilus 6

**S2 Supporting molecular dynamics information** **10**
Contact analysis in the 100 ns equilibrium simulations 20

**S3 Detailed molecular dynamics simulation methods** **24**
Preparation for steered molecular dynamics (sMD) simulations 24
sMD simulations 26
Extended equilibrium simulations 27
Steered Molecular Dynamics Movie Captions 27

**References** **28**

**S0 Optical tweezers instrumentation and sample preparation**

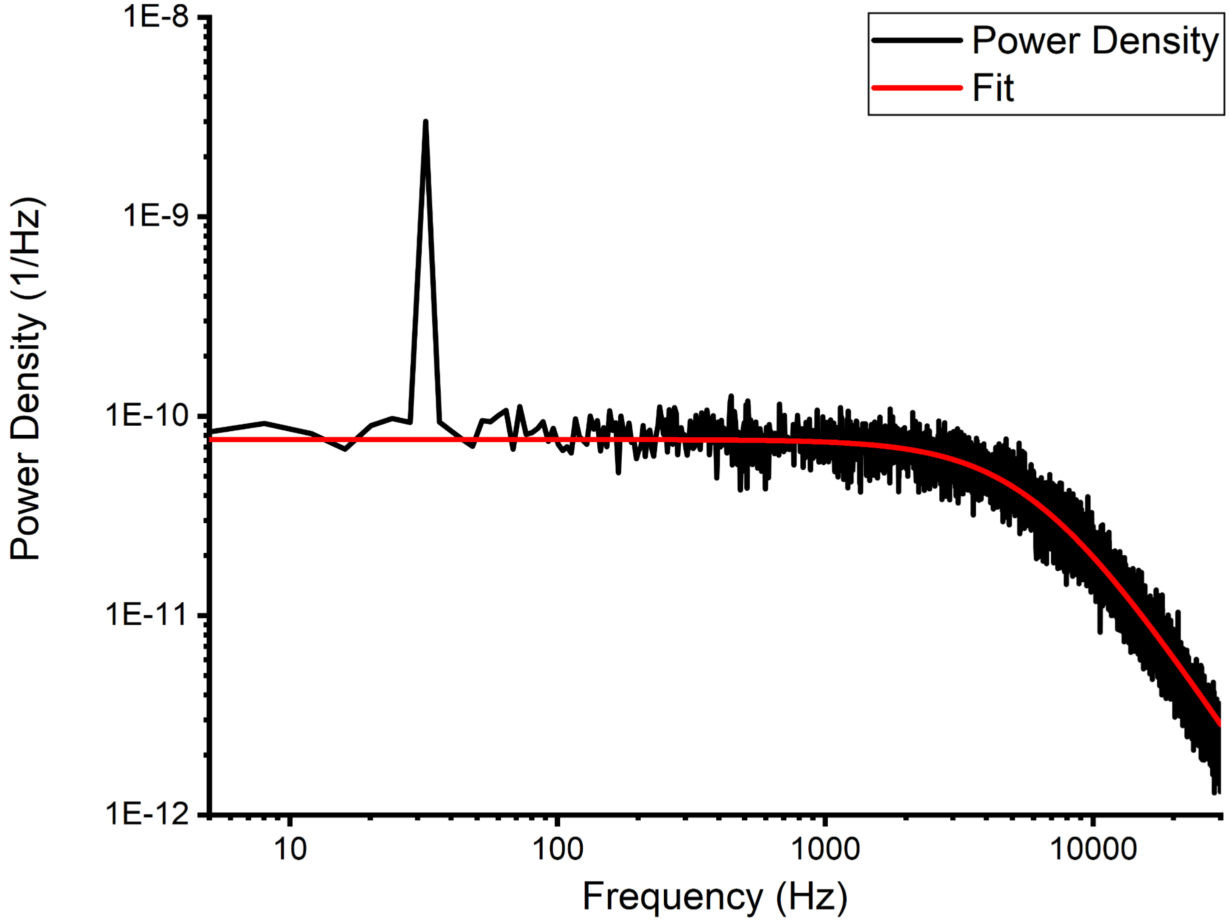


**Figure S1.** Example of an experimental power spectrum of a trapped bead (black). We have fitted the power spectrum with a Lorenztian model (red) given by $y = A/(f^2 + f_c^2)$ where $f$ is the frequency, $f_c$ the corner frequency and $A$ a fitting parameter related to the diffusion coefficient of the trapped bead and sensitivity of the detection system. The resulting fitting parameters are $A = 0.002663$, and $f_c = 5907$ Hz. The trapped bead is oscillated at 32 Hz, whose response is clearly visible in the data as a sharp peak in the power spectrum.

The optical tweezers instrumentation stands in a temperature controlled room on an actively isolated optical table (TMC). To reduce electrical and acoustic noise in the setup, we place computers and controllers in a separate room. We form the trap using a 1064 nm DPSS laser (Cobolt Rumba, Cobolt AB, Solna, Sweden) with a maximum power of 2 W. The laser beam is expanded to fill the numerical aperture of our microscope objective and introduced into the microscope using a dichroic mirror (DMSP650, Thorlabs, Newton, NJ). The laser light is then transmitted by the sample, scattered by the trapped object, and collected by the condenser. The collected light then interferes in the back focal plane of the condenser, which we image onto a 2D position-sensitive detector (PSD, 2L10YAG SU65 SPC02, Sitek Electro Optics, Sweden). The PSD tracks the centroid of the interference pattern and converts it to a voltage. We then filter the voltage with a programmable low pass anti-aliasing filter (LTC1064-2, Analog Devices, Wilmington, MA) before measuring it using a computer equipped with a data acquisition ( DAQ) card. We then process the collected data using LabVIEW.

To prepare a sample, we first suspend bacteria in 1xPBS to a concentration of 1:1000 from $OD_{600}$ = 1. Then we also suspend 1.04 μm polystyrene microspheres (4010A, Thermo Fisher Scientific, Waltham, MA) in 1xPBS. These microspheres are later used as force probes. To create a stable mounting point for bacteria, we prepare a 1:500 suspension of 9.5 um carboxylate-modified latex microspheres (product no. 2-10000, Interfacial Dynamics, Portland, OR) in Milli-Q water. 10 μl of this bead suspension is then dropped on a 24 x 60 mm coverslip (no. 1, Paul Marienfeld GmbH, Lauda-Königshofen, Germany). To fix these beads on the surface of the coverslip, we dry them in an oven for 60 min at 60 C. Then, we add a solution of 20 μl of 0.01 % poly-L-lysine (catalog no. P4832, Sigma-Aldrich, St. Louis, MO) to the coverslip, which we dry for 45 min at 37 C. This layer of poly-L-lysine helps the bacteria stay firmly attached during the measurement process.

To construct a sample chamber, we first add two strips of double-sided tape (Scotch, product no. 34-8509-3289-7, 3M) to one of the bead-coated coverslips. Then we place a 20 x 20 mm coverslip (no.1, Paul Marienfeld GmbH, Lauda-Königshofen, Germany) on top of the tape. This process forms a small gap between the lower and upper coverslips, and the tape acts as a spacer and a sealant. After that, we let capillary forces draw in 2 μl of suspended bacteria and 10 μl of suspended beads through the open sides of the chamber. We then seal these open sides using vacuum grease (Dow Corning, Midland, MI). Finally, we mount the completed sample chamber in a sample holder fixed to a piezo-stage (Physik Instrument, P-561.3CD stage) in the OT instrumentation.

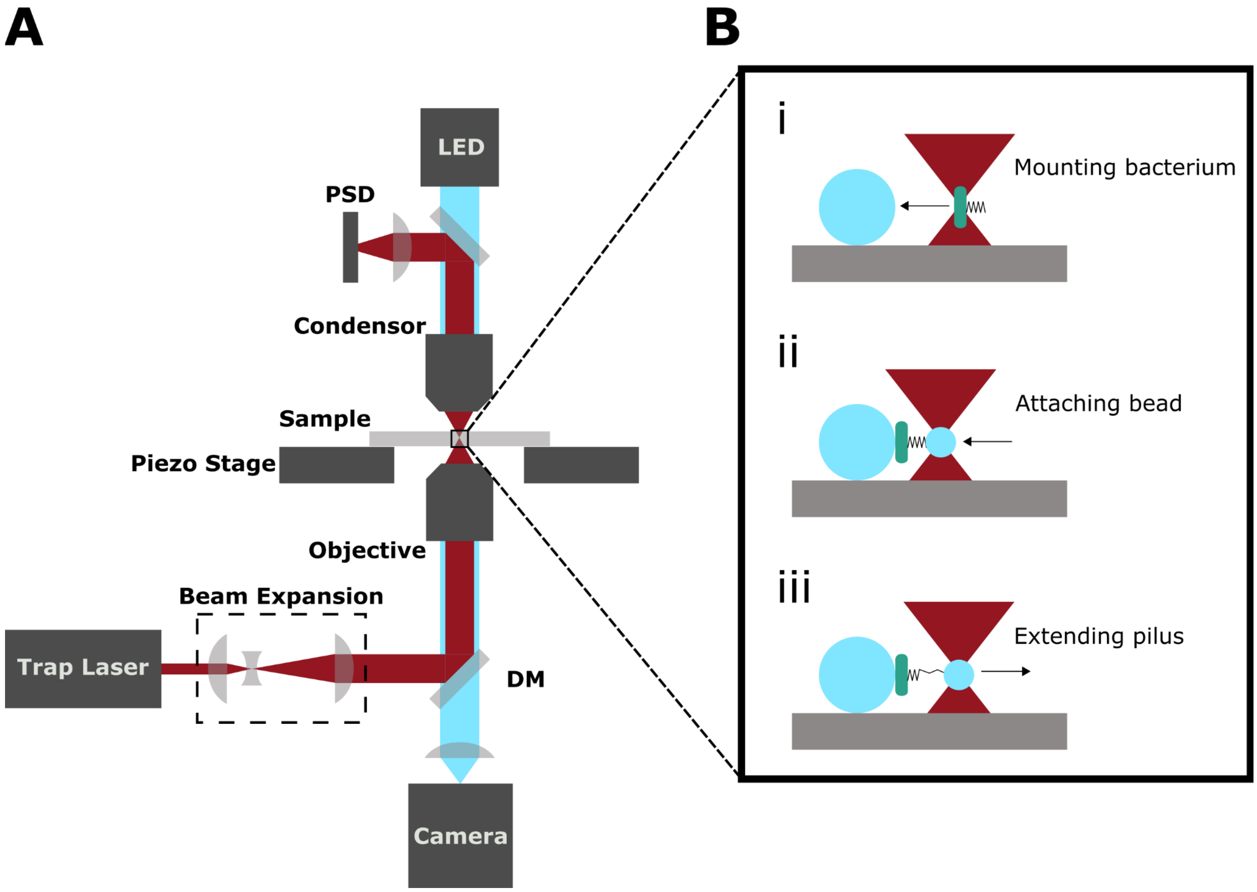


**Figure S2.** Schematic of the optical tweezer setup used to measure the coiling and uncoiling of pili. (A) We introduce the trap laser beam into the microscope using a dichroic mirror (DM). The laser beam is then focused by the objective into the sample to form the trap. To measure the position of a trapped microsphere we collect the light scattered by the sphere and transmitted through the sample with the condenser. The collected light interferes in the back focal plane of the condenser which we image on a 2D PSD. The PSD measures the centroid of the interference pattern and converts it to a voltage that we measure using a computer. (B) An illustration of a force-extension measurement. i) We trap a single bacterium and mount it onto a large microsphere that is fixed to the bottom coverslip of the sample chamber. ii) We trap a small microsphere that we attach to a pilus on the bacterium. iii) We separate the bacterium and trapped microsphere to apply a tensile force to the pilus and extend it partially. Then we track how the coiling and uncoiling of the pilus displaces the bead in the trap versus time.

### S1 Supporting force spectroscopy information and pilus modeling

#### *Measuring stiffness of an uncoiled pilus*

Due to the inherent force clamping of an unwinding pilus, we cannot easily assess its stiffness directly. Therefore, to measure the stiffness of a partially unwound pilus, we use the equipartition theorem. This method allows us to assess the stiffness of the harmonic potential formed by both the trap and the pilus. For small displacements, as in our setup, we can assume that both the trap and the pilus are harmonic. Thus, we can estimate the total stiffness of trap and pilus, $k_{tot}$, using the equipartition theorem defined as,

$$k_{tot} = \frac{k_B T}{<x^2>}, \quad (1)$$

where $k_B$ is Boltzmann's constant, $T$ is the temperature, and $<x^2>$ is the variance of the trapped bead's position. As the pilus and optical trap are connected to the bead in parallel, their stiffnesses should be additive [1]. Thus, we can estimate the stiffness of the pilus, $k_p$, using our measured value for $k_{tot}$ and $k_t$ using

$$k_p = k_{tot} - k_t. \quad (2)$$

To estimate the stiffness of a pilus from our data, we used a moving variance with a window length of 3000 samples. We converted this moving variance to stiffness using Eq. 1 and 2 and created a histogram of the values. From this histogram, we estimated the most likely stiffness by taking the stiffness with the most counts. We show an example of two histograms like this for a pilus extended 3 μm and 8 μm into region II, Figure S3. This figure shows that the measured stiffness decreases as the pilus gets extended, and the histogram approaches a normal distribution. The asymmetry of the stiffer grey histogram comes from the contribution of the binding and unbinding events, which create high variance spikes in the signal. This effect disappears for the less stiff red histogram because when the stiffness goes down, we lose temporal and spatial resolution and do not resolve these binding and unbinding events clearly.

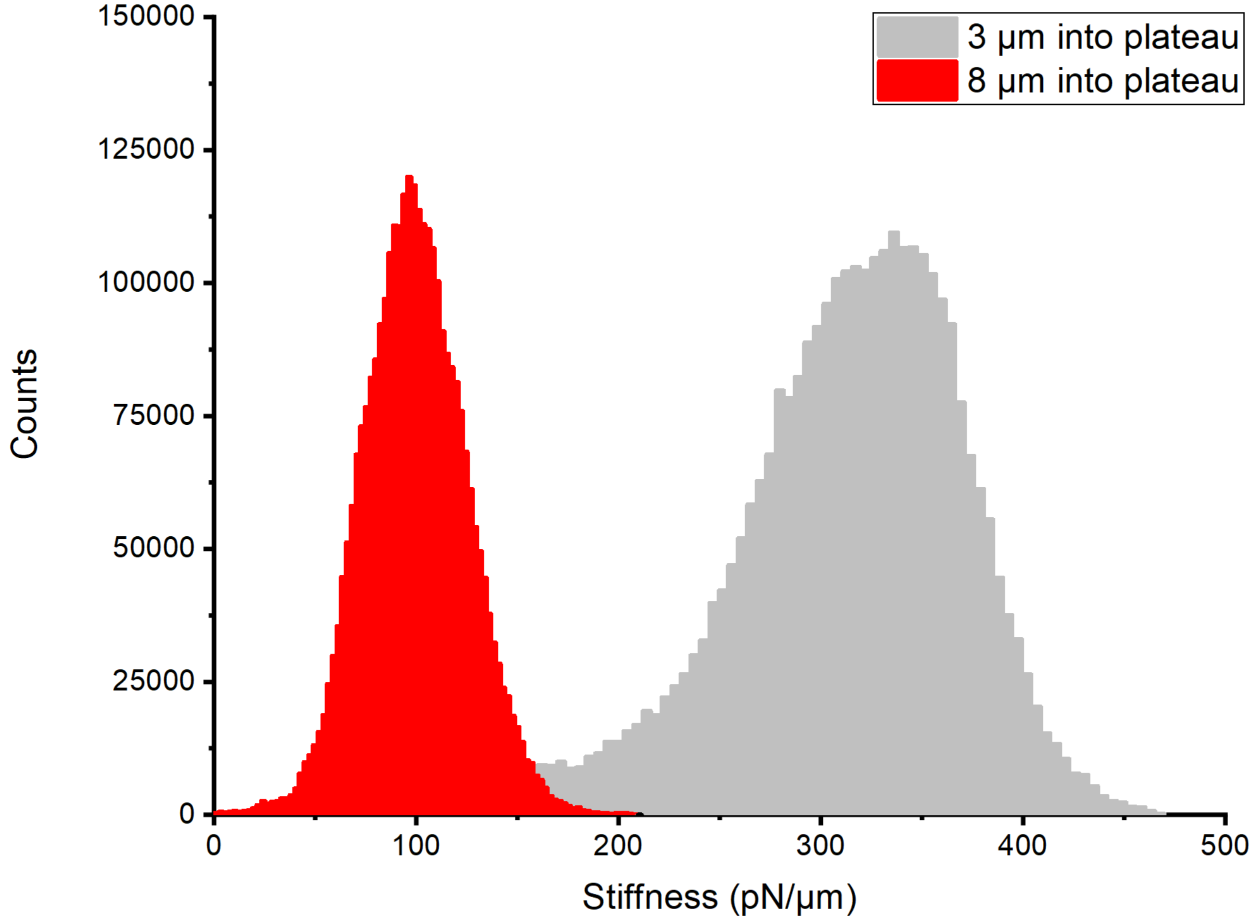


**Figure S3.** Stiffness of pilus measured using a moving variance histogram of the force data with a window length of 3000 samples for two different extensions of the same pilus.

***Compliance correction***

When measuring the length changes of a pilus when a subunit loses and regains contact with its neighbors using an optical tweezer, in which the trap's position is held constant, we need to account for the fact that the force applied to the pilus does not stay constant as it shortens or elongates. This changing force will change the relative degree of extension of the pilus at the same time as it elongates or shortens. This effect causes our measured displacement of the bead to be less than the actual length change of the pilus [1]. The relationship between the displacement of the bead, $x_m$, and the length change of the pilus, $\Delta R$ , is given by

$$x_m = \Delta R \frac{k_t + k_p}{k_p}. \tag{3}$$

***Stiffness model for an uncoiling pilus***

We model an uncoiling pilus as a combination of an extensible worm like chain (WLC) [2], representing the coiled rod, and a freely jointed chain (FJC) [3], representing the uncoiled linear region. For the uncoiled region the FJC end-to-end distance, $R_u$, versus force, $F$, relationship is given by

$$R_u = L_u(coth(\frac{FL_k}{k_B T}) - \frac{k_B T}{Fl_k}), \quad (5)$$

where $T$ is the temperature, $k_B$ is Boltzmann's constant, $L_u$ is the contour length, and $l_k$ is the Kuhn length. Here, the end-to-end distance $R_u$ is the straight line between the first and last subunit, whereas the contour length $L_u$ is the length of the backbone of all subunits in the uncoiled configuration, as diagrammed in Figure S4.

For the coiled region, the WLC end-to-end distance, $R_c$, versus force relationship is given by

$$R_c = L_c(1 - \frac{1}{2}(\frac{k_B T}{FP_c})^{1/2} + \frac{F}{K_c}), \quad (4)$$

where, $L_c$ is the contour length, $P_c$ is the persistence length, and $K_c$ is the enthalpic stiffness of the coiled pilus. Thus, the total end-to-end distance $R_{tot}$ vs force for the pilus, if we assume that the coiled and uncoiled regions are experiencing the same force, is given by

$$R_{tot} = R_u + R_c. \quad (6)$$

To model the uncoiling of the pilus we treat it as being composed of $N_{tot}$ subunits where $N_b$ is the number of subunits in the bound state that forms the coiled region of the pilus and $N_u$ is the number in the unbound state, forming the uncoiled region, and $N_{tot} = N_c + N_u$. Each subunit in these two states contributes to the total contour length of the pilus with the lengths $x_b$ and $x_u$ for the bound and unbound states, respectively. Thus, the total contour length can be calculated by

$$L_{tot} = L_u + L_c = N_b x_b + N_u x_u. \quad (7)$$

During uncoiling, the pilus will start with all subunits in the bound state, so $N_b$=$N_{tot}$, and transition to a fully uncoiled structure with $N_u$=$N_{tot}$. This uncoiling will change the contour lengths of the coiled and uncoiled regions of the structure, leading to an end-to-end distance versus force relationship that changes with the degree of uncoiling. By combining these equations, we can express the total end-to-end distance versus force as a function of the number of unbound subunits in the following way

$$R_{tot}(F,\ N_u) \ = \ N_u x_u (coth(\frac{Fl_k}{k_B T}) - \frac{k_B T}{Fl_k}) + (N_{tot} - N_u) x_c (1 - \frac{1}{2}(\frac{k_B T}{FP_c})^{1/2} + \frac{F}{K_c}). \tag{8}$$

From this equation, we can relate the change in end-to-end distance, $\Delta R_{tot}$, when a subunit unbinds, to the actual change in total contour length, $\Delta L_{tot}$, by the following equations

$$\Delta R_{tot} = R_{tot}(F, N_u) - R_{tot}(F, N_u + 1) = x_b(1 - \frac{1}{2}(\frac{k_B T}{Fc})^{1/2} + \frac{F}{K_c}) - x_u(coth(\frac{Fl_k}{k_B T}) - \frac{k_B T}{Fl_k}), \tag{9}$$

$$\Delta L_{tot} = x_u - x_b. \tag{10}$$

Further, by taking the inverse of $\frac{dR_{tot}(F, N_u)}{dF}$ we can get the stiffness, $k_{pili}$, of a partially uncoiled pilus as

$$k_{pili} = (\frac{dR_{tot}(F, N_u)}{dF})^{-1} = (x_u N_u (\frac{k_b T}{F^2 l_k} - csch^2(\frac{Fl_k}{k_b T})) \ + \ x_b(N_{tot} - N_u)(\frac{1}{4}(\frac{k_b T}{FP_c})^{-1/2} \frac{k_b T}{F^2 P_c} + \frac{1}{K_c}))^{-1}. \tag{11}$$

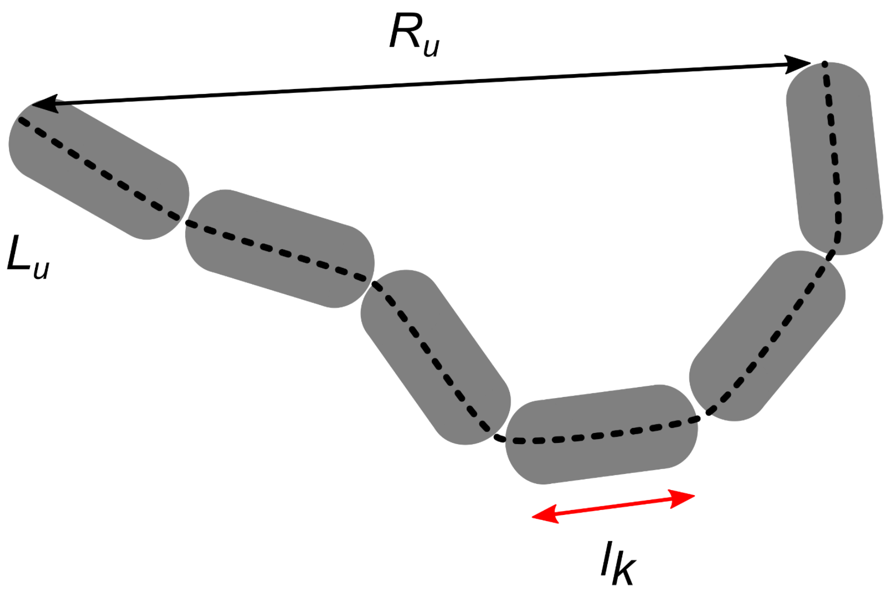


**Figure S4** Schematic showing the difference between end-to-end distance $R_u$(solid line), and contour length $L_u$(dashed line) of a polymer assembled from subunits with a Kuhn length $l_k$(red line).

To validate this model, we measured the stiffness of P pili (measure on 3 individual pili) for various degrees of uncoiling using the method outlined below. Comparing these data to the calculated stiffness using our model with parameters $P_c$ = 1 µm, $K_c$ = 1300 pN, $N_{tot}$ = 2000, $l_k$ = 5.1 nm, $x_b$ = 0.754 nm, $F$ = 28 pN, and $x_u$ = 5.1 nm, shows good agreement, Figure S5. As the value $l_k$ represents the length of a link in the FJC, the fact that our result closely matches the length of a PapA subunit, ~5 nm, lends credence to our model. Further, if we compare this to a previously used stiffness model for P pili [4], we see that this older model did not reflect our measured stiffness versus elongation behavior and underestimated the pilus stiffness. This indicates that it is more suitable to model the uncoiling pilus as a WLC connected in series with a FJC.

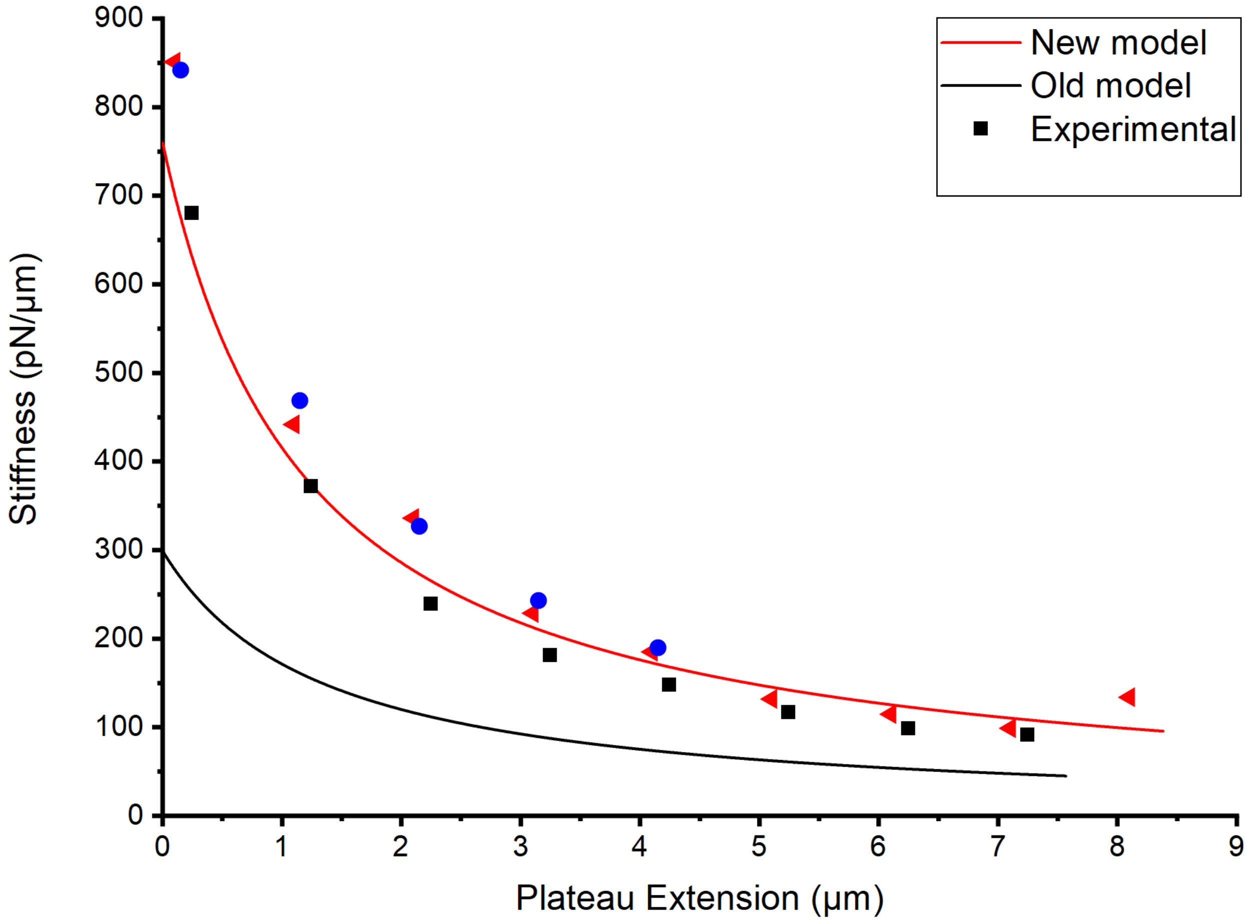


**Figure S5.** A comparison of our stiffness model (red line), previous stiffness model (black line), and three experimental data sets represented by the red triangles, blue dots and black squares.

### S2 Supporting molecular dynamics information

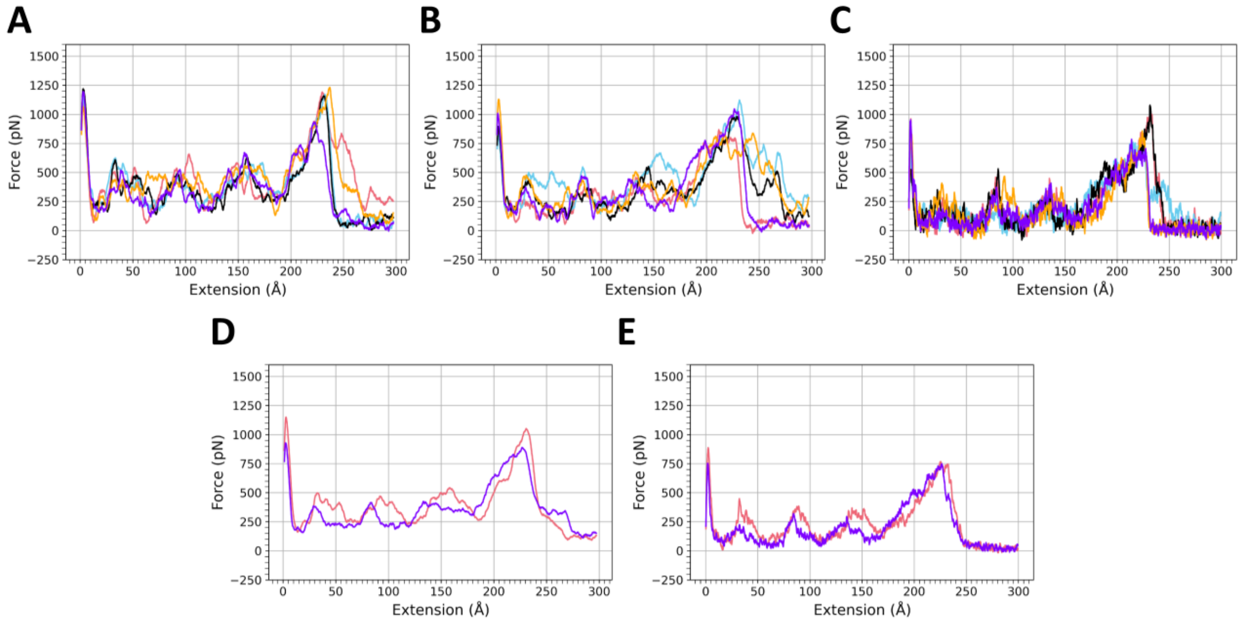


**Figure S6.** Force versus extension curves for all five of the v = 5 Å/ns steered molecular dynamics simulations with (A) the staple residues present, and (B) the staple residues removed. (C) Force versus extension curves for all five of the v = 1 Å/ns steered molecular dynamics simulations with the staple residues removed. The corresponding figure with the staple residues present for the v = 1 Å/ns steered molecular dynamics simulations is Figure 3 of the main text. In panels (A), (B) and (C) pink = run 1, blue = run 2, black = run 3, orange = run 4, and purple = run 5. (D) Force versus extension curves for the v = 5 Å/ns steered molecular dynamics simulations averaged over the five separate runs for the system including the staple (pink) and with the staple residues removed (purple). (E) Force versus extension curves for the v = 1 Å/ns steered molecular dynamics simulations averaged over the five separate runs for the system including the staple (pink) and with the staple residues removed (purple). The force curve data are a 1 ns running average.

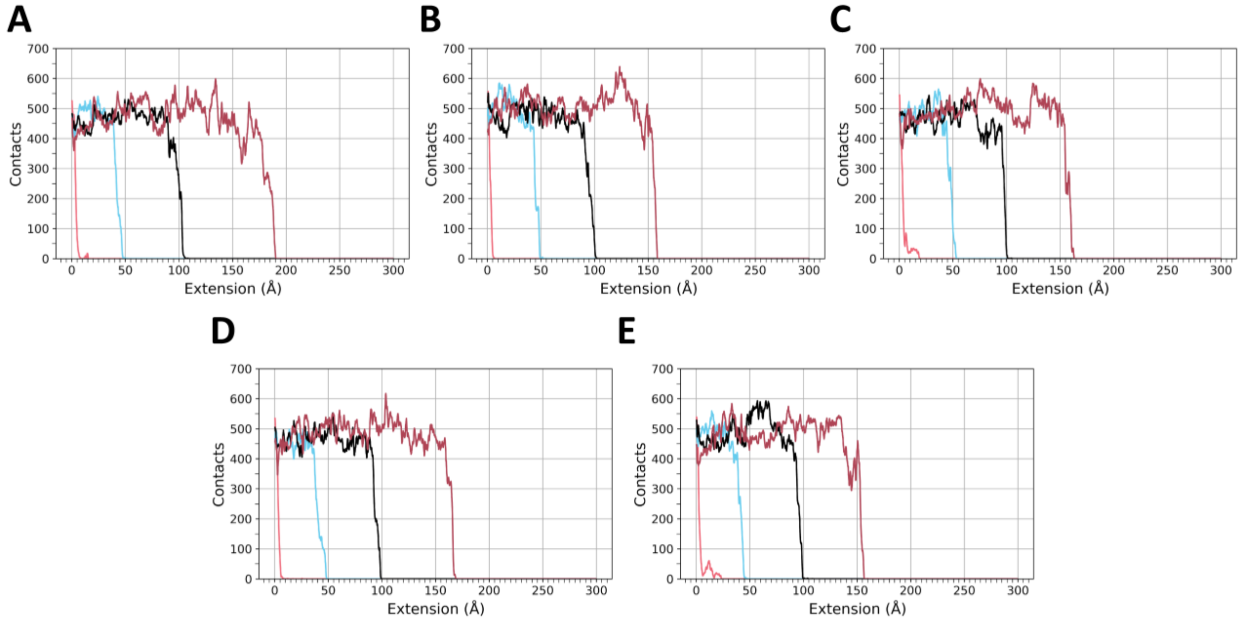


**Figure S7.** Total number of contacts between subunits 1 and 4 (pink), 2 and 5 (blue), 3 and 6 (black), and 4 and 7 (dark red), for the v = 1 Å/ns simulations (A) run 1, (B) run 2, (C) run 3, (D) run 4, and (E) run 5. Data are plotted as a function of extension for ease of comparison to the force vs extension curves. The contacts data are a 1 ns running average. Data are for the simulations with the staple.

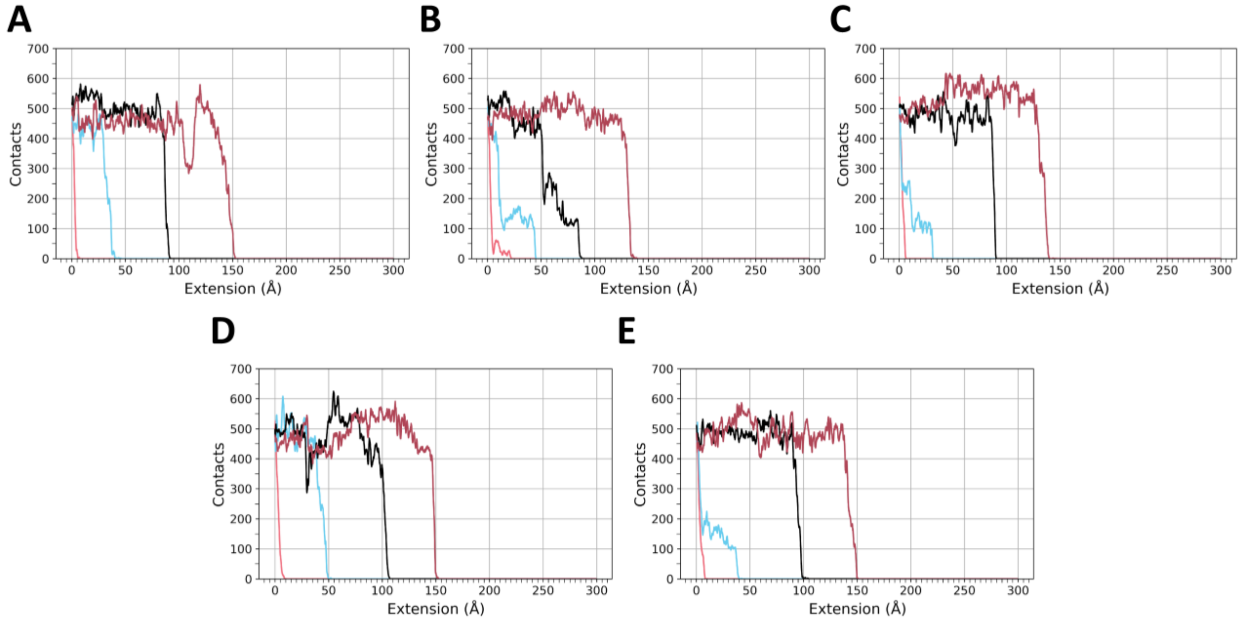


**Figure S8.** Total number of contacts between subunits 1 and 4 (pink), 2 and 5 (blue), 3 and 6 (black), and 4 and 7 (dark red), for the v = 1 Å/ns simulations (A) run 1, (B) run 2, (C) run 3, (D) run 4, and (E) run 5. Data are plotted as a function of extension for ease of comparison to the force vs extension curves. The contacts data are a 1 ns running average. Data are for the simulations without the staple.

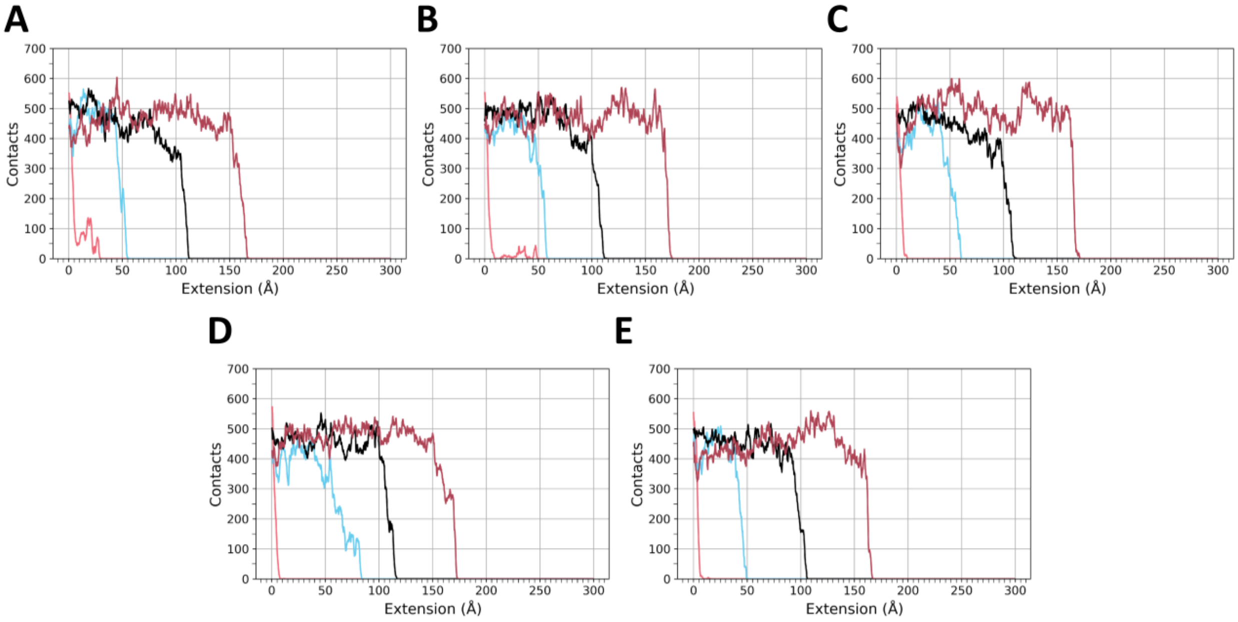


**Figure S9.** Total number of contacts between subunits 1 and 4 (pink), 2 and 5 (blue), 3 and 6 (black), and 4 and 7 (dark red), for the v = 5 Å/ns simulations (A) run 1, (B) run 2, (C) run 3, (D) run 4, and (E) run 5. Data are plotted as a function of extension for ease of comparison to the force vs extension curves. The contacts data are a 200 ps running average. Data are for the simulations with the staple.

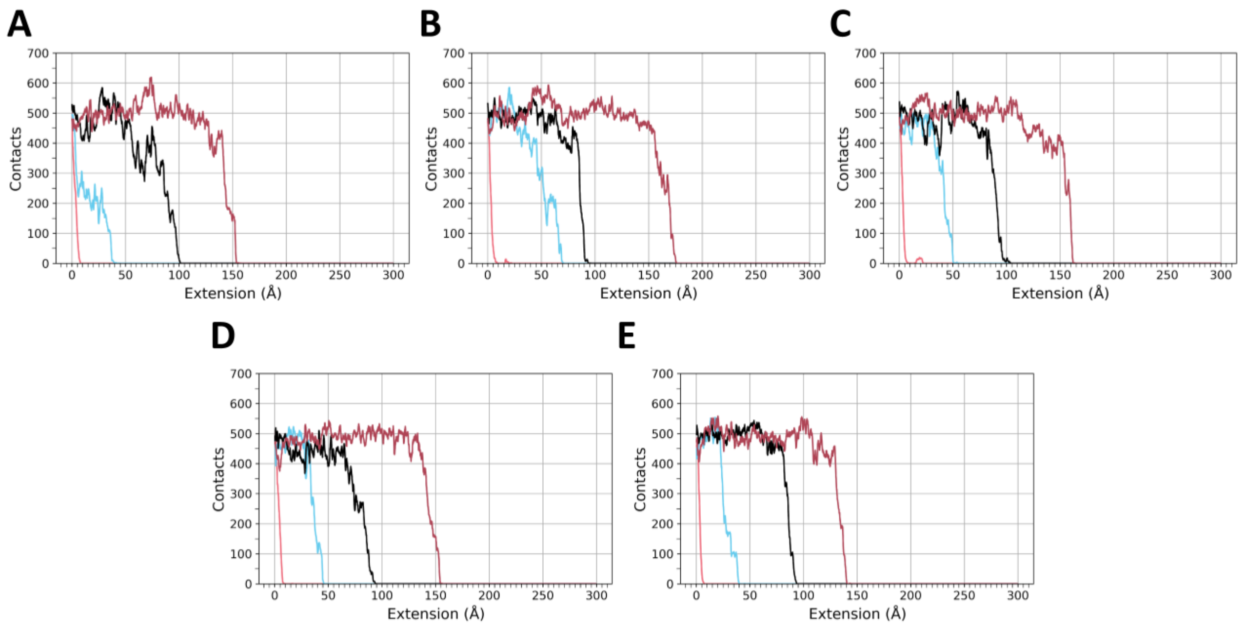


**Figure S10.** Total number of contacts between subunits 1 and 4 (pink), 2 and 5 (blue), 3 and 6 (black), and 4 and 7 (dark red), for the v = 5 Å/ns simulations (A) run 1, (B) run 2, (C) run 3, (D) run 4, and (E) run 5. Data are plotted as a function of extension for ease of comparison to the force vs extension curves. The contacts data are a 200 ps running average. Data are for the simulations without the staple.

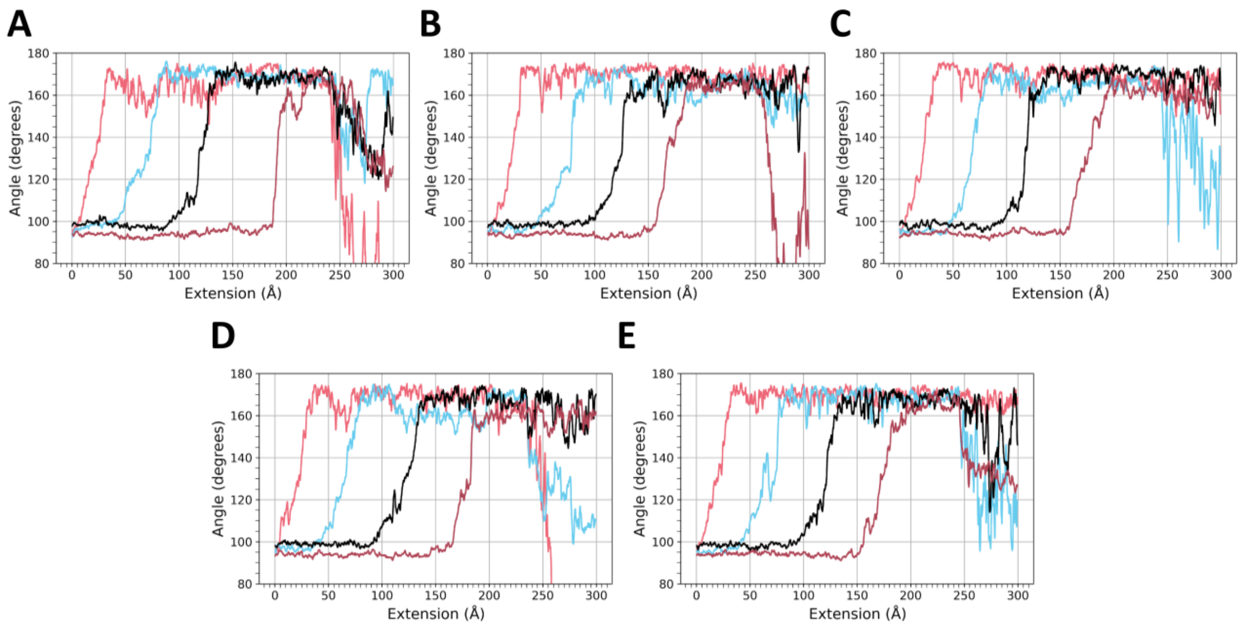


**Figure S11.** Angle between residues Gly7, Val18, and Cys22 (location of residues can be viewed in the main manuscript as Figure 3E, inset) for the v = 1 Å/ns simulations. As in the main manuscript Figure 3, the four curves correspond to the alignment of subunit 1 (pink curve), subunit 2 (blue curve), subunit 3 (black curve) and subunit 4 (dark red curve) with the filament axis. When the angle is 90 degrees, the subunit is perpendicular to the filament axis and when the angle is 180 degrees, the subunit is aligned with the filament axis. The angle data are a 1 ns running average. The panels correspond to (A) run 1, (B) run 2, (C) run 3, (D) run 4, and (E) run 5. Data are for the simulations with the staple.

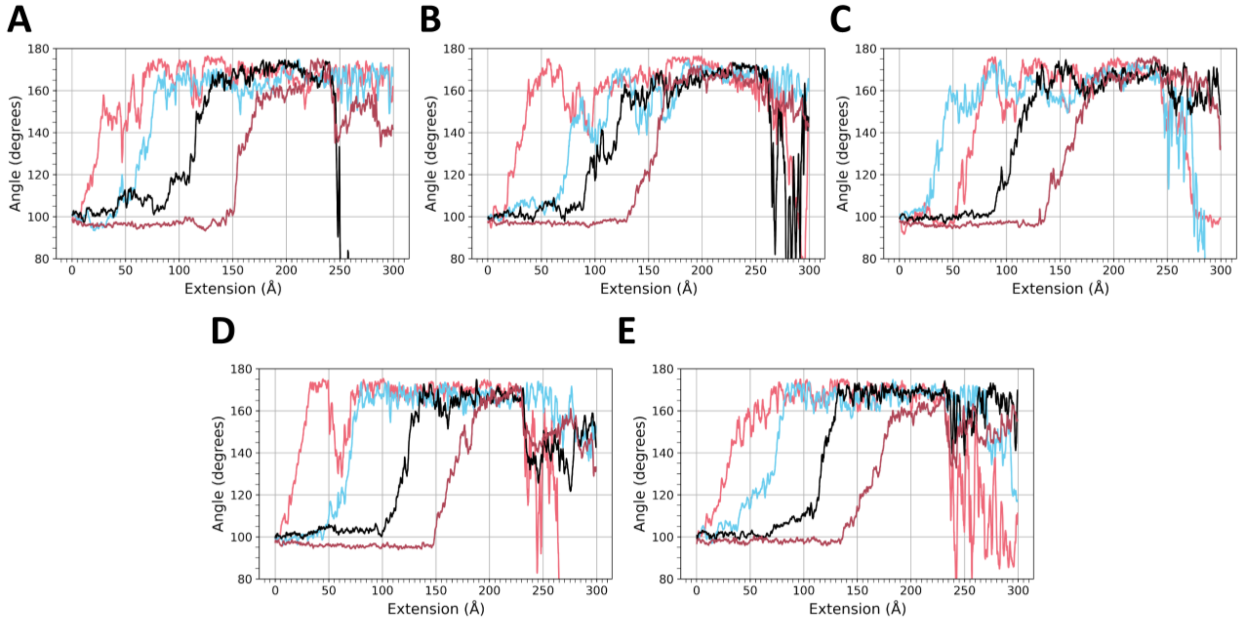


**Figure S12.** Angle between residues Gly7, Val18, and Cys22 (location of residues can be viewed in the main manuscript in the inset image of Figure 3E) for the v = 1 Å/ns simulations. As in the main manuscript Figure 3, the four curves correspond to the alignment of subunit 1 (pink curve), subunit 2 (blue curve), subunit 3 (black curve) and subunit 4 (dark red curve) with the filament axis. When the angle is 90 degrees, the subunit is perpendicular to the filament axis and when the angle is 180 degrees, the subunit is aligned with the filament axis. The angle data are a 1 ns running average. The panels correspond to (A) run 1, (B) run 2, (C) run 3, (D) run 4, and (E) run 5. Data are for the simulations without the staple.

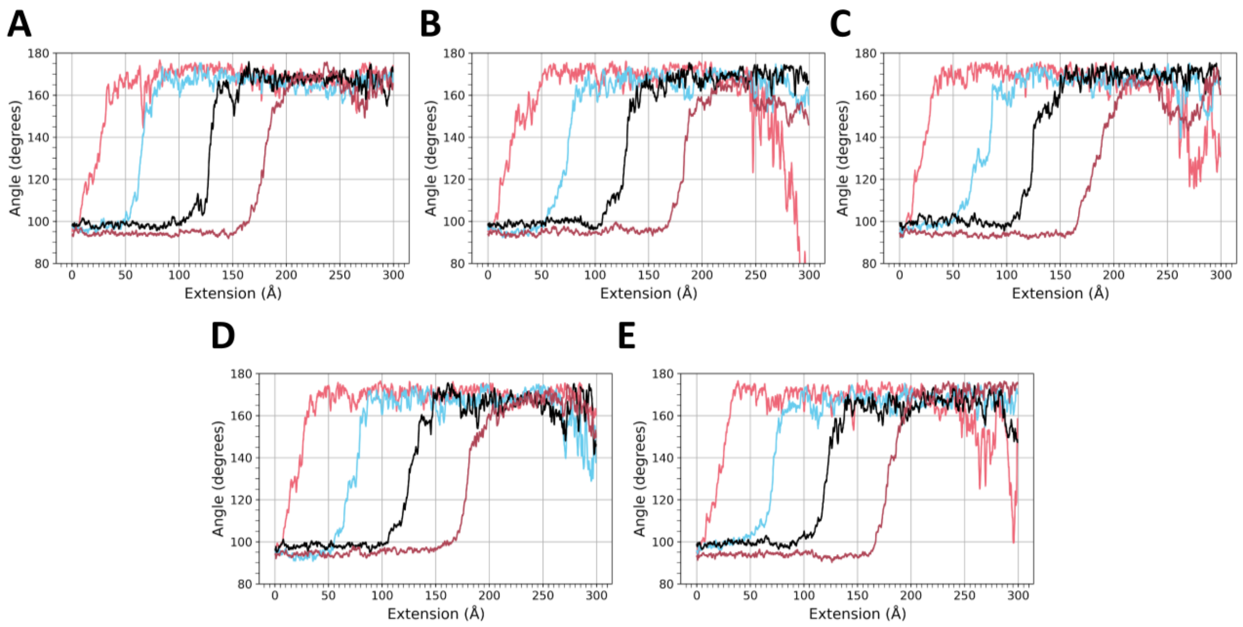


**Figure S13.** Angle between residues Gly7, Val18, and Cys22 (location of residues can be viewed in the main manuscript in the inset image of Figure 3E) for the v = 5 Å/ns simulations. As in the main manuscript Figure 3, the four curves correspond to the alignment of subunit 1 (pink curve), subunit 2 (blue curve), subunit 3 (black curve) and subunit 4 (dark red curve) with the filament axis. When the angle is 90 degrees, the subunit is perpendicular to the filament axis and when the angle is 180 degrees, the subunit is aligned with the filament axis. The angle data are a 200 ps running average. The panels correspond to (A) run 1, (B) run 2, (C) run 3, (D) run 4, and (E) run 5. Data are for the simulations with the staple.

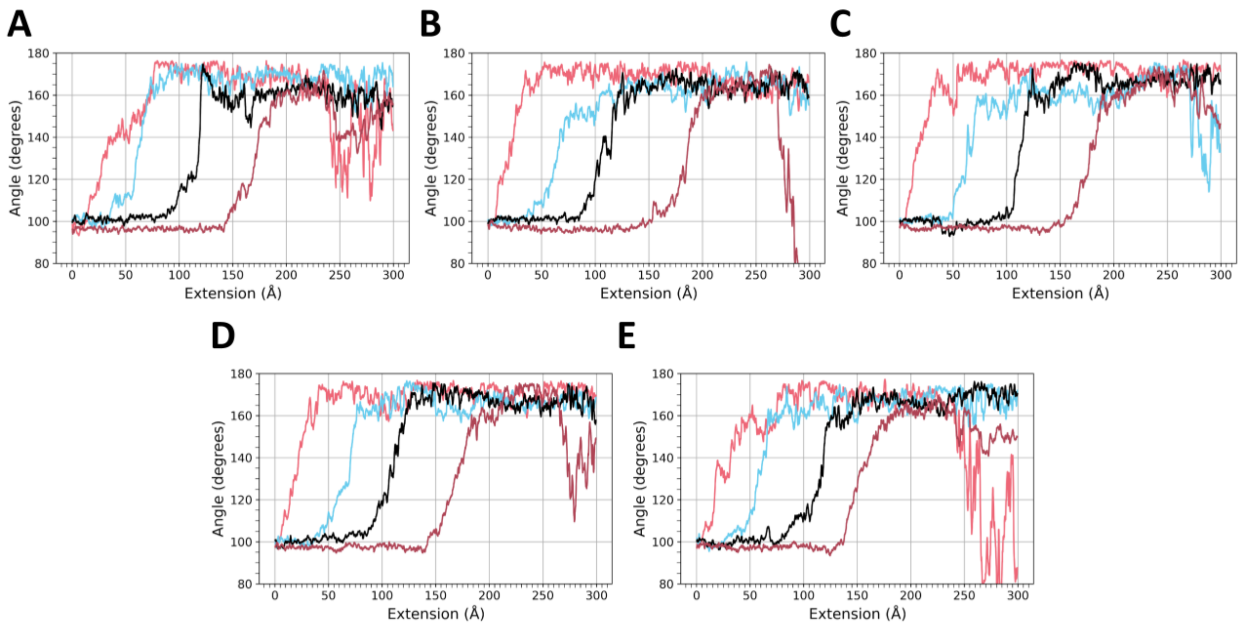


**Figure S14.** Angle between residues Gly7, Val18, and Cys22 (location of residues can be viewed in the main manuscript in the inset image of Figure 3 in panel E) for the v = 5 Å/ns simulations. As in the main manuscript Figure 3, the four curves correspond to the alignment of subunit 1 (pink curve), subunit 2 (blue curve), subunit 3 (black curve) and subunit 4 (dark red curve) with the filament axis. When the angle is 90 degrees, the subunit is perpendicular to the filament axis and when the angle is 180 degrees, the subunit is aligned with the filament axis. The angle data are a 200 ps running average. The panels correspond to (A) run 1, (B) run 2, (C) run 3, (D) run 4, and (E) run 5. Data are for the simulations without the staple.

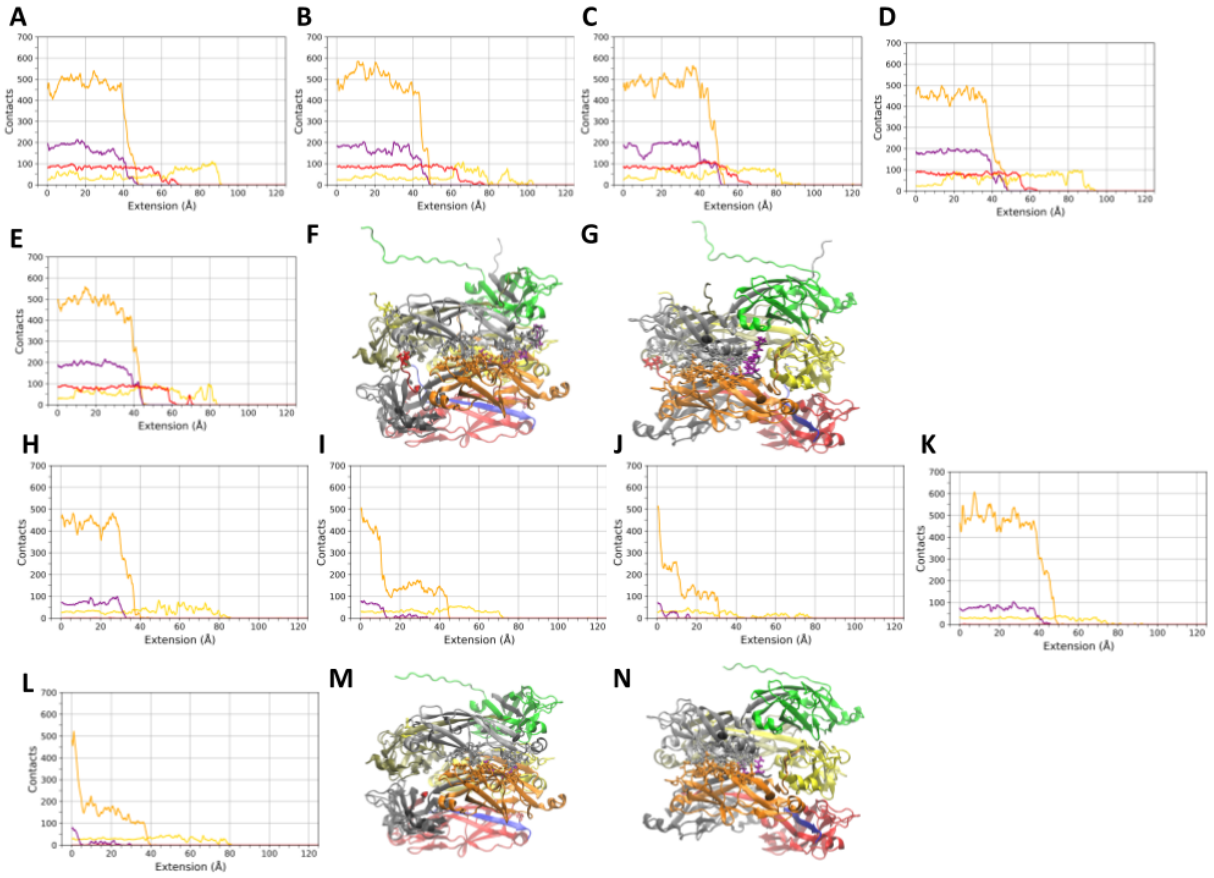


**Figure S15.** Total number of contacts between subunits 2 and 4 (yellow), 2 and 5 (orange), 2 and 6 (purple), and 2 and 7 (red), for the v = 1 Å/ns simulations (A) run 1, (B) run 2, (C) run 3, (D) run 4, and (E) run 5. Data are plotted as a function of extension for ease of comparison to the force vs extension curves. The plot is cut off at 125 Å since contacts between the depicted regions have all dropped to zero at that point. The contacts data are a 1 ns running average. Data in (A) - (E) are for the simulations with the staple. (F) and (G) show ribbon diagram images of the 7mer with groups of atoms that are initially within 4 Å of the light grey subunit and the yellow, orange, dark grey, and red subunits depicted in stick view. Colors of the lines in panels A through E are the same as the subunit coloring using the same color scheme as in panels F and G, except the contacts between the light grey subunit and the dark grey subunit are shown in purple for clarity. For the simulations without the staple, the total number of contacts between subunits 2 and 4 (yellow), 2 and 5 (orange), 2 and 6 (grey), and 2 and 7 (red), are shown for the v = 1 Å/ns simulations (H) run 1, (I) run 2, (J) run 3, (K) run 4, and (L) run 5. (M) and (N) show images of the 7mer without the staple with groups of atoms initially within 4 Å of the light grey subunit and the yellow, orange, dark grey, and red subunits depicted. Colors of the lines in panels (H) - (L) are the same as the subunit coloring in panels M and N, except the contacts between the light grey subunit and the dark grey subunit are shown in purple for clarity.

#### *Contact analysis in the 100 ns equilibrium simulations*

For the purposes of the data in the tables S1-S4, the presence of a contact between two residues in a frame of a trajectory is defined to be any two non-hydrogen atoms of the two residues coming within a 4 Å cutoff distance of one another. The presence of contacts was calculated at 500 ps intervals throughout the 100 ns trajectories. The time fraction is the ratio of the number of analyzed trajectory frames in which contacts are present divided by the total number of analyzed trajectory frames. Subunit-subunit contacts that occur for at least ⅓ of the equilibrium simulation time are reported. An asterisk symbol next to an "interacting residue" indicates that this interaction was also described by Hospenthal *et al.* in Figure S3 of [14].

**Table S1. Contact analysis from equilibrium simulations - subunits 2 and 7**
Steered molecular dynamics showing subunit-subunit contacts that occur at least ⅓ of the equilibrium simulation time between subunit 2 and subunit 7 of the 7mer filament.

| Residue (Subunit 2) | Interacting residue | Interacting Subunit | Time fraction bond is present (with staple) | Time fraction bond is present (no staple) |
|---|---|---|---|---|
| Phe42 | Pro2 * | 7 | 0.85 | Not present |
| Ala45 | Ala1 * | 7 | 0.71 | Not present |
| | Pro2 | 7 | 0.93 | Not present |

**Table S2. Contact analysis from equilibrium simulations - subunits 2 and 5**
Steered molecular dynamics showing subunit-subunit contacts that occur at least ⅓ of the equilibrium simulation time between subunit 2 and subunit 5 of the 7mer filament.

| Residue (Subunit 2) | Interacting residue | Interacting Subunit | Time fraction bond is present (with staple) | Time fraction bond is present (no staple) |
|---|---|---|---|---|
| Gln106 | Asn96 * | 5 | 1.00 | 0.95 |
| | Gly97 | 5 | 0.86 | 0.98 |
| | Gly98 * | 5 | 0.62 | 0.86 |
| Ala108 | Pro84 | 5 | 0.62 | 0.69 |
| | Asp94 * | 5 | 0.98 | 0.69 |
| | Thr95 | 5 | 1.00 | 0.10 |
| | Asn96 | 5 | 0.74 | 0.96 |
| Gly109 | Asp94 * | 5 | 0.74 | 0.57 |
| Asn122 | Gly83 | 5 | 0.58 | 0.29 |
| | Pro84 * | 5 | 0.73 | 0.46 |
| Thr123 | Thr82 * | 5 | 0.88 | 0.9 |
| | Asp115 | 5 | 0.61 | 0.62 |
| Lys125 | Thr82 | 5 | 0.82 | 0.69 |
| | Phe158 | 5 | 0.98 | 0.94 |
| | Asn159 * | 5 | 1.00 | 0.82 |
| Asp126 | Asn159 | 5 | 1.00 | 0.53 |
| Asn129 | Asn157 | 5 | 0.34 | 0.50 |
| Val130 | Val155 * | 5 | 0.37 | 0.39 |
| | Asn157 * | 5 | 0.97 | 0.99 |
| His132 | Asn96 * | 5 | 0.81 | 0.96 |
| | Ser153 * | 5 | 0.96 | 0.99 |
| | Ala154 | 5 | 0.55 | 0.61 |
| | Val155 * | 5 | 0.78 | 0.90 |
| Tyr133 | Asn96 | 5 | 0.97 | 0.75 |
| Thr134 | Asn96 * | 5 | 1.00 | 0.91 |
| | Ser153 | 5 | 0.42 | 0.60 |

**Table S3. Contact analysis from equilibrium simulations - subunits 2 and 6**
Steered molecular dynamics showing subunit-subunit contacts that occur at least ⅓ of the equilibrium simulation time between subunit 2 and subunit 6 of the 7mer filament.

| Residue (Subunit 2) | Interacting residue | Interacting Subunit | Time fraction bond is present (with staple) | Time fraction bond is present (no staple) |
|---|---|---|---|---|
| Asp62 | Pro2 | 6 | 0.93 | Not present |
| | Thr3 * | 6 | 0.98 | Not present |
| Thr64 | Thr3 | 6 | 0.59 | Not present |
| Lys125 | Gln8 * | 6 | 1.00 | 0.99 |
| Asp126 | Thr3 | 6 | 0.52 | Not present |
| | Pro5 * | 6 | 0.68 | Not present |
| | Gln8 | 6 | 0.42 | 0.41 |
| Gly127 | Pro5 | 6 | 0.59 | Not present |
| | Gln8 | 6 | 0.96 | 0.66 |
| Glu128 | Gln8 * | 6 | 1.00 | 0.99 |
| Val130 | Lys10 | 6 | 0.35 | 0.55 |

**Table S4. Contact analysis from equilibrium simulations only found in no staple system - subunits 2 and 5**
Steered molecular dynamics showing new subunit-subunit contacts that occur at least ⅓ of the equilibrium simulation time between subunit 2 and subunit 5 of the 7mer filament only in the system with no staple.

| Residue (Subunit 2) | Interacting residue | Interacting Subunit | Time fraction bond is present (with staple) | Time fraction bond is present (no staple) |
|---|---|---|---|---|
| Asp53 | Ser153 | 5 | Not present | 0.34 |
| Lys67 | Asp115 | 5 | Not present | 0.39 |
| | Gly116 | 5 | Not present | 0.33 |
| Gly109 | Asn96 | 5 | Not present | 0.91 |

### S3 Detailed molecular dynamics simulation methods

#### *Preparation for steered molecular dynamics (sMD) simulations*

All simulated systems were prepared using the program tLeap which is included with Amber20/AmberTools21 [5]. In the initial structure file, the 7mer pilus filament was aligned so that the helical axis was along the z-direction. This orientation was used for all simulations. Note that for the subunit at the “base” of the filament, the inserted β strand from the prior subunit is also included in the simulated structure (using the first 20 amino acids of the N-terminal extension (NTE)). Protein parameters were described by the FF14SB force field [6] and the parameters for water molecules were described by the TIP3P force field [7]. Monovalent counterions were described using the parameters of Joung and Cheatham [8].

The 7mer systems were solvated in a rectangular water box. In the x and y directions (perpendicular to the filament axis) a 12 Å buffer to the periodic cell edge was used, and in the z-direction (parallel to the filament axis) a 160 Å buffer was implemented. This allowed for the filament to be extended along the filament axis during sMD simulations while maintaining its solvation. Overall charge neutralization was achieved by adding 35 $Na^+$ counterions to the system.

Preparation of the systems for sMD simulations used an approach very similar to that in simulations of the 3mer system [9], but modified accordingly to simulate the larger 7mer system for the current study. Differences in protocol between simulations with and without the staple are pointed out specifically in the text below. Note that for simulations without the staple, the first five amino acids were removed from the N-terminal end of each subunit in the 7mer system, so that the first amino acid in each subunit becomes Gln6. Energy minimization was accomplished using 3000 steps of steepest descent and 2000 steps of conjugate gradient while applying a restraint force constant of 10.0 kcal $mol^{-1}$ $Å^{-2}$ to the alpha carbons. Subsequently, system heating was carried out using two separate stages. First, the system temperature was increased from 0 K to 100 K in the NVT ensemble over 20 ps, and then the temperature was held at 100 K for 30 ps. Second, heating using the NPT ensemble was performed and the system temperature was increased from 100 K to 300 K (20 ps duration) and then held at a temperature of 300 K (80 ps duration). During the heating stages, a restraint force constant of 10.0 kcal $mol^{-1}$ $Å^{-2}$ was applied to the alpha carbons. After heating, equilibration of the systems was carried out over seven stages while maintaining a temperature of 300 K in the NPT ensemble, and using a changing set of atom restraints depending on the equilibration stage. Table S5 below describes how the restraints were applied during the equilibration stages of simulation. After the equilibration was completed, the last frame was used as the initial coordinates for the sMD simulations.

**Table S5. Restraints during equilibration stages to prepare for sMD simulations**

| Stage | Duration (ps) | Restrained selection | Restraint strength (kcal $mol^{-1}$ $Å^{-2}$) |
|---|---|---|---|
| 1 | 200 | All alpha carbons | 10.0 |
| 2 | 200 | All alpha carbons | 5.0 |
| 3 | 200 | All alpha carbons | 2.5 |
| 4 | 200 | All alpha carbons | 0.5 |
| 5 | 200 | All alpha carbons | 0.1 |
| 6 | 1000 | Alpha carbons of Subunit 7 (and its inserted NTE β-strand from Subunit 8), Subunit 6, Subunit 5 (except for its NTE β-strand) | 0.1 |
| 7 | 4000 | Alpha carbons of Subunit 7 (and its inserted NTE β-strand from Subunit 8), Subunit 6, Subunit 5 (except for its NTE β-strand), Subunit 1 | 0.1 |

***sMD simulations***

To implement the sMD protocol, the jar = 1 [10] flag in Amber20 was set and all of the sMD simulations were carried out using the NPT ensemble with a temperature of 300 K. The sMD simulations require the definition of a collective variable (CV) used to describe the extension of the system. We used the z-distance between the center of mass of the alpha carbons of subunits 7 (and its inserted NTE β-strand from Subunit 8), subunit 6, and subunit 5 (except for its NTE β-strand), and the alpha carbons of the terminal ($1^{st}$) subunit in the 7mer (except for the alpha carbons in its NTE β-strand to allow that region to remain freely mobile during 7mer extension). We refer to those selections as the “fixed” and “pulled” selections, respectively. In order to apply the sMD force along the z-direction, the AMBER20 fxyz option was used. The fixed selection was restrained by applying a 0.5 kcal $mol^{-1}$ $Å^{-2}$ restraint to the alpha carbons in that atom group. This allowed us to have the bottom three subunits mimic a segment of the filament base being adhered to a surface as the top subunit is pulled, allowing four out of the seven subunits to extend away from the base in the sMD simulations. The restraints applied to the three subunits at the base also eliminated the possibility for overall rotations and translations of the system during the sMD simulations. Systems were extended by approximately 300 Å along the z-direction, and this led to overall elongation of the 7mer system, and in some cases eventually led to breakage of the system between subunits (.e.g, separation by the extraction of an NTE β-strand). The spring constant parameter for the pulling spring in the AMBER input files was set to a value of 5.0, which results in a pulling spring stiffness of 10.0 kcal $mol^{-1}$ $Å^{-2}$. The force and amount of extension were saved every 2 ps. During the heating, equilibration, and sMD simulations the integration timestep was set to 2 fs, and the SHAKE algorithm was applied to hydrogen bonds [11]. The Langevin thermostat was used for temperature control, and we implemented a collision frequency of 1 $ps^{-1}$. During stages of the simulation that were carried out in the NPT ensemble, the Monte Carlo barostat [12] was used to maintain a pressure of 1 atm. All stages of simulation employed a real space interaction cutoff of 8 Å, and long range electrostatics were handled using the particle mesh Ewald method [13].

***Extended equilibrium simulations***

We also carried out two extended equilibrium simulations of the 7mer filaments, one for the system with the staple residues, and one for the simulation of the system without the staple residues. The equilibrium simulations lasted for 100 ns. They were carried out in the NPT ensemble, again using the Langevin thermostat and the Monte Carlo barostat as described above to maintain a temperature of 300 K and a pressure of 1 atm. During these simulations, restraints were only applied to the alpha carbons of the bottom three subunits of each system as described in Table S5 above, with the following modification: the first 8 amino acids of the subunit 8 NTE β-strand, as well as the first 8 amino acids of subunits 7 and 6, were allowed to remain unrestrained (without the staple this was residues 6-8, as 1-5 were deleted). Neither long equilibrium simulation had restraints applied to the terminal (1st) subunit in the 7mer filament. The restraints applied to the alpha carbon atoms of the bottom three subunits prevented overall translation and rotation of the 7mer during the extended equilibrium simulations.

***Steered Molecular Dynamics Movie Captions***

**Movie S1.** Movie generated in VMD of the v = 1 Å/ns steered molecular dynamics simulations of the P pilus system with the staple for runs 1 through run 5.

**Movie S2.** Movie generated in VMD of the v = 1 Å/ns steered molecular dynamics simulations of the P pilus system without the staple for runs 1 through run 5.

**Movie S3.** Movie generated in VMD of the v = 5 Å/ns steered molecular dynamics simulations of the P pilus system with the staple for runs 1 through run 5.

**Movie S4.** Movie generated in VMD of the v = 5 Å/ns steered molecular dynamics simulations of the P pilus system without the staple for runs 1 through run 5.
